# The resource-rational dynamics of evidence accumulation

**DOI:** 10.64898/2026.04.15.718716

**Authors:** Mengting Fang, Jiang Mao, Tobias H. Donner, Alan A. Stocker

## Abstract

Evidence accumulation is a fundamental aspect of human decision-making. However, how the precise temporal structure of evidence shapes the accumulation process has not been systematically studied. As a result, current understanding of evidence accumulation remains largely limited to its time-averaged behavior. We tested human subjects in a visual estimation task in which they inferred the angular position of an unknown source from a noisy stimulus sequence. Introducing systematic temporal perturbations, i.e., breaks of different durations and at different positions in the otherwise regular evidence sequence, revealed that subjects actively compensated for the memory loss endured during the break by dynamically enhancing evidence integration and memory maintenance immediately after the break. We derived a new time-continuous Bayesian updating model that is dynamically constrained by optimal performance-effort trade-offs. With two free parameters determining the overall resource-efficiencies of encoding and memory maintenance, the model accurately predicts the rich dependencies of subjects’ accumulation behavior on the evidence schedule, including subjects’ individual tendencies to emphasize either early (primacy) or late (recency) samples in the evidence sequence. Our results suggest that evidence accumulation is a non-stationary, dynamically controlled process that optimally balances the information gained from incoming evidence against the cognitive effort required to acquire and maintain it. The proposed model is general and should apply broadly across many task domains.

## Introduction

Evidence accumulation is the process by which an observer updates and maintains beliefs about the state of the world as new evidence becomes available. Efficient and accurate evidence accumulation is crucial for making accurate decisions, and hence important from an evolutionary standpoint^1,2,3,4^. The human brain can accumulate evidence over a wide range of time-scales, from the order of less than a second, as in determining the direction of motion of a random-dot kinematogram^5,6^, to years and entire lifespans, as in the formation of consumer brand preferences based on infrequent product experiences^7,8,9,10^. Independent of these large scale differences, evidence accumulation relies on two general mechanisms: evidence must be registered and used to update beliefs when available (encoding), and beliefs must be kept in memory throughout the accumulation phase (maintenance). Both mechanisms are subject to noise and other resource limitations, which, in combination with task specific statistics and reward objectives, determine accumulation behavior^11,12,13,14,15,16^. However, previous studies almost exclusively examined evidence accumulation under conditions in which the sensory evidence rate (evidence per unit time) was constant; even so sensory evidence rarely arrives with regular rates under real-life conditions. In the few studies that used irregular evidence streams^12,17,18,19^, the irregularities were random yet, on average, the rate remained constant. Constant (or averaged constant) evidence rates mask the full complexity of the underlying dynamics. As a result, previous studies may have missed important dynamic aspects of evidence accumulation, and thus have painted a rather simplistic picture of its computational complexity.

With this study we systematically examined how the precise schedule of a sensory evidence sequence affects the accumulation process. In a series of psychophysical experiments, we asked human subjects to estimate an unknown continuous variable (angular position) based on a sequence of visual stimuli representing noisy samples of the unknown position. We systematically created perturbations in the evidence stream by introducing a break of varying duration or position in the otherwise regular sample sequence. Across all conditions, we found that the first sample after the break contributed significantly stronger to subjects’ estimates than the same sample in the same sequence but without a break. This increase in weight, which we refer to as “peak-after-break” effect, was consistent across all subjects regardless of their individual tendencies toward primacy or recency. The effect cannot be explained by any previous models of evidence accumulation for continuous estimation tasks (e.g., bump-attractor models^16^). Thus, we derived a new, time-continuous resource-rational model of evidence accumulation. The model assumes that evidence is accumulated through Bayesian updating. However, both encoding of incoming sensory evidence and memory maintenance of the accumulated belief are formulated as resource constrained, dynamic processes that continuously maintain an optimal balance between expected task performance and the required resource cost. With two free parameters, expressing the relative resource efficiencies of encoding and memory maintenance, our model accurately predicts the observed dynamics of sequential evidence integration under tested conditions with varying break durations and positions. It effectively captures the peak-after-break effect as well as the large range of weighting patterns across individual subjects (primacy and recency) observed in our data. Our findings suggest that perceptual evidence accumulation is an actively controlled, resource-rational process with non-stationary dynamics that reflect a continuously negotiated, optimal trade-off between performance and effort.

## Results

We conducted three psychophysical experiments that shared the same basic design (Fig. 1A). Subjects were presented with a rapid sequence of eight visual stimuli representing noisy samples from a source with an unknown angular position. After stimulus presentation, subjects had to provide an estimate of the source’s position by adjusting a cursor, upon which they immediately received feedback about the accuracy of their estimate by showing the actual position of the source. The three experiments were designed to systematically characterize how a break in the evidence stream affected evidence accumulation in human subjects. By analyzing the effect of a single temporal perturbation per trial, and probing subjects evidence accumulation characteristics with a fine-grained estimation task, we were able to quantitatively characterize and, ultimately, model the actively controlled information processes that guide human sensory evidence accumulation.

**Figure 1:**
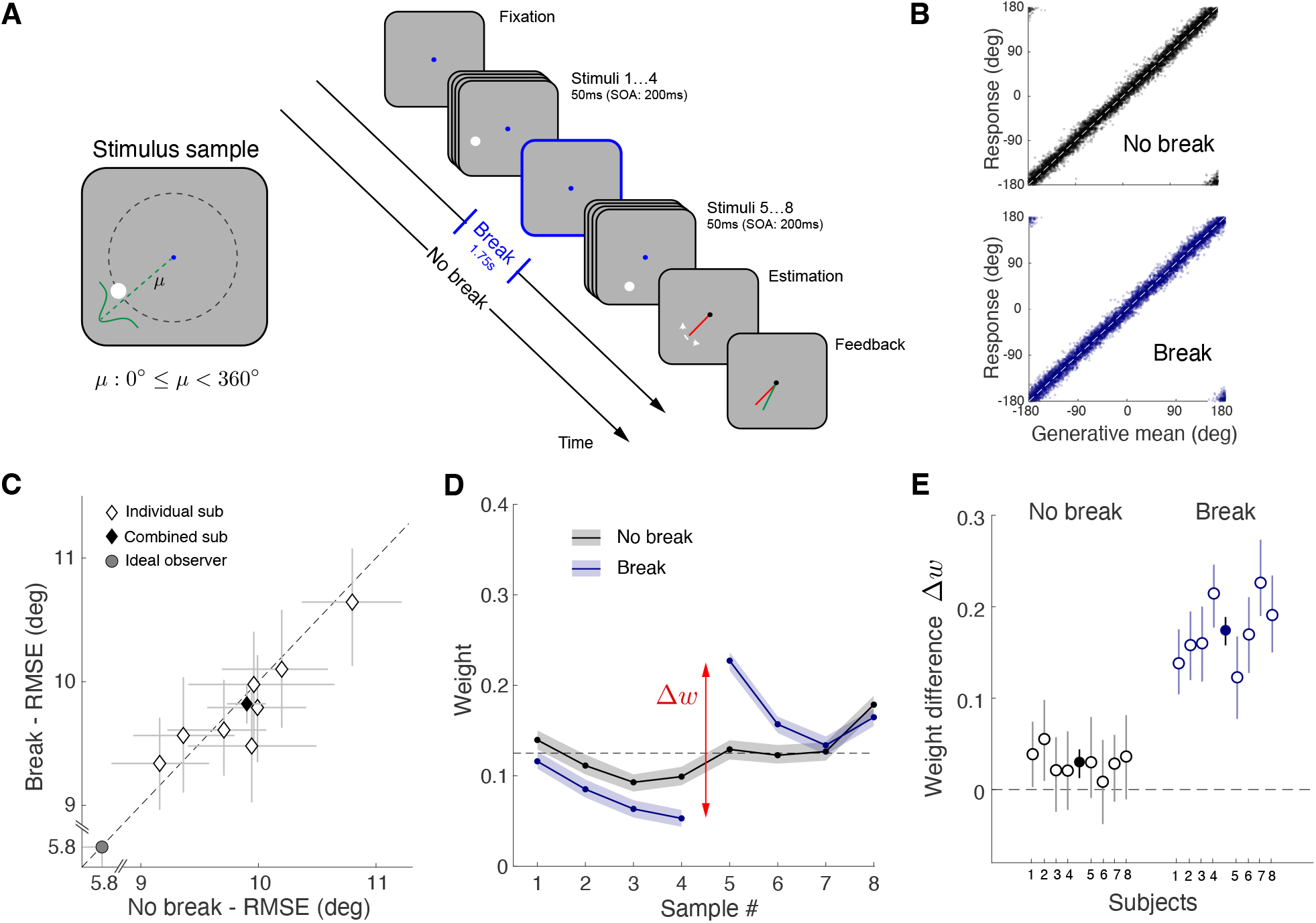
Sequential evidence accumulation for a continuous variable. (A) Experiment 1: Subjects (N=8) were instructed to estimate the angular position of an unknown generative mean (i.e., the angular position of the unknown source) based on 8 sequentially presented noisy samples (white dots on gray background). Samples were drawn from a Gaussian with fixed variance centered at the generative mean *µ*. Samples were regularly presented (“No break” condition) or with an interruption of 1.75 s after the 4th sample (“Break” condition). Sample sequences of every trial were random and different for individual subjects, but identical base sequences were used for both conditions. (B) Response distributions as a function of generative mean (combined subject). (C) Estimation performance measured as root mean squared error (RMSE). (D) Temporal weighting kernels obtained from a circular regression a nalysis. Kernels consistently exhibited a pronounced “peak-after-break” (i.e., a disproportionately large, positive weight difference Δ*w* between the 5th and the 4th sample) in the “Break” condition (combined subject; see Fig. 4E for kernels of individual subjects). (E) This effect was observed for all subjects (combined subject: filled circles). Shaded areas and error bars in all panels represent the 95% confidence intervals computed over 200 bootstrapped samples of the data. Supplementary Figure S1: Comparison of interleaved and non-interleaved conditions. Supplementary Figure S2A: Error-distribution relative to generative mean for individual subjects.

### Experiment 1: A break in the evidence stream causes active compensation in evidence encoding

Experiment 1 aimed to test the general impact of a temporal perturbation in an otherwise regular evidence stream. After training, the subjects performed the basic task (Fig. 1A) in three sessions. In the first session, samples were presented without interruption (“No break” condition). In the second session, the sequence was always interrupted after the fourth sample for a fixed duration (“Break” condition). In the third session, both conditions were randomly interleaved. We generated a set of 550 unique base sample sequences for each subject, which were used in both the “No break” and “Break” conditions across all three sessions, totaling 2200 trials. Thus, our comparison of evidence accumulation across conditions is based on identical stimulus sequences, differing only by a uniform random rotation applied to each sequence. In the following, we combined the data for each condition from the non-interleaved and interleaved sessions because subjects’ behavior did not show any significant difference across these sessions other than a minimal performance increase due to learning (see Supplementary Fig. S1).

Subjects performed well under both conditions. The estimates were very similarly distributed, closely following the true generative mean (Fig. 1B). While estimation accuracy differed between subjects, every subject performed at near identical levels under the two conditions (Fig. 1C). With subjects’ performances substantially lower than ideal observer performance, evidence accumulation and maintenance were clearly impaired by sensory processing noise and other inefficiencies. Given that memory leakage is one known inefficiency in evidence maintenance^20,21,22,23^, we would have expected a worse performance under the “Break” condition given the extra leak during the relatively long break duration. A circular regression analysis provided some insight into why performance did not significantly drop. The resulting temporal weighting profiles for the “No break” and “Break” trials revealed substantial differences (Fig. 1D). For “No break” trials the weights show an overall rather continuous and approximately uniform pattern. For “Break” trials, however, the weight of the first sample after the break, and weaker the second, is significantly higher compared to that of the last sample before the break. This effect is consistent across all subjects regardless of their individual tendencies towards primacy, recency, or uniform temporal integration (Fig. 1E; see Supplementary Fig. S1 for individual subjects).

Note that any leaky integrator model would qualitatively exhibit a peak-after-break effect simply because some of the information about the first four samples is lost during the break. However, an adaptive encoding mechanism is required in order to compensate for the loss during the second half of the sample sequence. Existing models do not have such mechanism and thus can not explain our experimental results (see also Supplementary Fig. S4). In the following, we introduce a new Bayesian model of evidence accumulation, in which dynamic mechanisms guide the precision of evidence encoding and maintenance. Our key assumption is that these mechanisms are controlled by a resource-rational process that dynamically maintains an optimal trade-off between performance and effort throughout the entire trial sequence.

### Dynamic model of resource-rational evidence accumulation

A Bayesian observer updates its belief about a world state (its probability) with each new piece of information according to Bayes’ rule (Fig. 2A). Applied to our experiment, we assume that the observer keeps a representation of the probability of the generative mean *µ* (i.e., the angular position of the unknown source) throughout the trial. The generative mean in the experiment was randomly sampled and thus has a uniform prior *P*_0_(*µ*) = 𝒰_[0,2*π*)_. Whenever a stimulus sample *S*_*i*_ is presented, the observer obtains a noisy measurement *θ*_*i*_ of the sample value, and updates its belief representation accordingly as

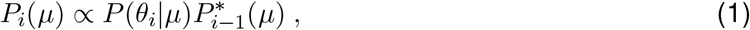

where 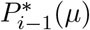 is the belief representation before the stimulus presentation, and

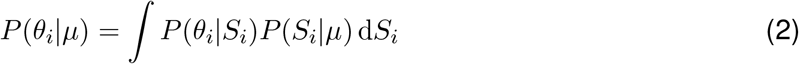

is the likelihood function; *P*(*θ*_*i*_|*S*_*i*_) represents the sensory encoding probability subject to sensory processing noise and *P*(*S*_*i*_|*µ*) the probability of the sample distribution for a given *µ*. During the inter-stimulus interval, we assume that the belief representation is subject to memory decay, changing from *P*_*i*_(*µ*) to 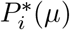 due to memory noise. At the end of the trial, the observer derives a final estimate 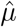 based on the posterior 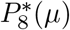 and a specific loss function. This is the general formulation of a Bayesian updating model where the sensory encoding noise and the memory noise are the two inefficiencies that account for a drop of performance below the level of the ideal observer. New in our model is that we assume that these two noise sources are dynamically controlled at any moment in time according to local objectives. Specifically, stimulus encoding and memory maintenance are governed by dynamic processes that ensure that cognitive effort is optimally balanced against expected performance. We refer to these processes as performance-effort trade-off (PET) modules.

**Figure 2:**
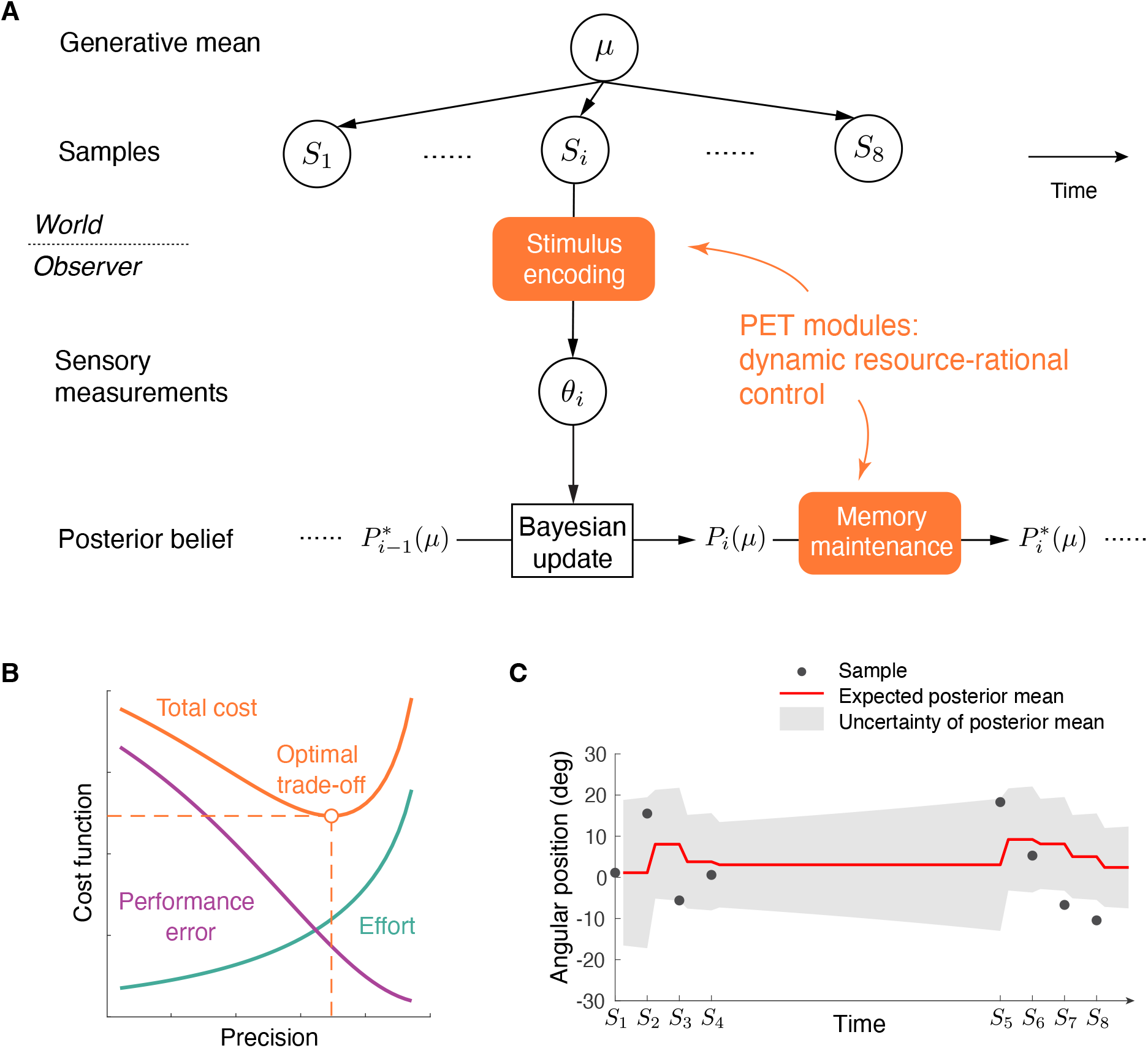
Dynamic evidence accumulation model. (A) During the presentation of each sample *S*_*i*_, the visual system registers (encodes) a noisy sensory measurement *θ*_*i*_ of the sample and then updates the current prior belief 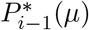 about the generative mean *µ* with the likelihood function based on the measurement. During the inter-stimulus interval, the observer maintains the representation of the updated belief with some memory noise. Importantly, we assume encoding and memory noise to be dynamically controlled by a performance-effort trade-off (PET) computation. (B) PET module: As the representational precision increases (less noise), the effort increases and the performance error decreases. Minimizing the weighted sum of the two cost functions determines the optimal precision (i.e., optimal noise level). (C) Example illustration of the evolution of the expected posterior mean (red line) across a sample sequence in the “Break” condition. Black dots represent the angular position of each sample, and the gray shaded area indicates the uncertainty of the posterior mean (± 1 SD).

Encoding of sample *S*_*i*_ results in a noisy measurement *θ*_*i*_, which is then used to update the belief about the generative mean *µ* (Fig. 2B). The PET module formulates this problem as minimizing a joint cost function consisting of an effort term and a performance inaccuracy term. Our formulation of this trade-off is inspired by the information bottleneck^24^. We express the encoding effort as mutual information between the sensory measurement *θ*_*i*_ and the stimulus sample *S*_*i*_

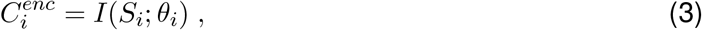

and the performance inaccuracy as the entropy of the generative mean after Bayesian updating

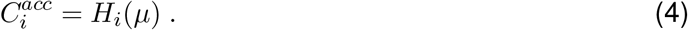

The optimal trade-off consists of choosing an encoding *P*(*θ*_*i*_|*S*_*i*_) that minimizes the weighted sum of the encoding cost and the inaccuracy,

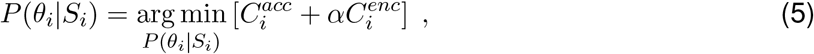

where *α* is the weight of the encoding cost relative to the inaccuracy (Fig. 2B). Larger values of *α* make encoding more costly, which results in lower optimal encoding precision; and vice versa.

A similar PET formulation governs memory maintenance. We assume that during the *i*th interstimulus interval, the belief *P*_*i*_(*µ*) randomly drifts and widens due to memory decay. Let 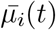 be the mean of *P*_*i*_(*µ*; *t*) at time *t* within the *i*th inter-stimulus interval. From time *t* to *t* + Δ*t*, the representation of *µ* drifts with 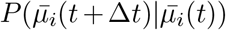 (memory noise). The lost information about 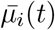 can be inferred retrospectively from 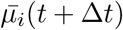, so the probability of *µ* at *t* + Δ*t* is

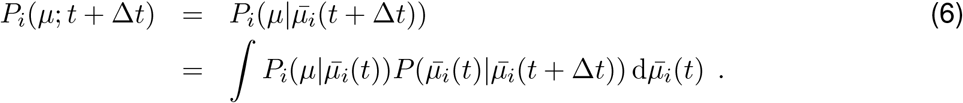

Because *µ* has a uniform prior, 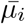 also has a uniform prior,

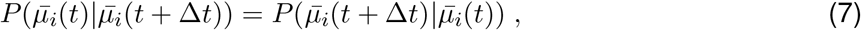

and we can compute *P*_*i*_(*µ*; *t* + Δ*t*) using Eq.(6) with 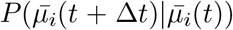 determined by the drift model. Again, we consider the maintenance trade-off as allowing 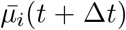 to be a noisy representation of 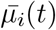 that preserves the optimal amount of information about *µ* for the invested maintenance effort. The corresponding PET module formulates this problem as minimizing a joint cost function consisting of an effort term for memory maintenance and a performance term for memory distortion. The memory maintenance effort for a time interval of Δ*t* is defined as the mutual information between the mean of *P*_*i*_(*µ*; *t*) and *P*_*i*_(*µ*; *t* + Δ*t*) maintained over Δ*t*,

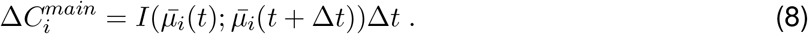

Memory distortion is defined as the increase in entropy of *µ*,

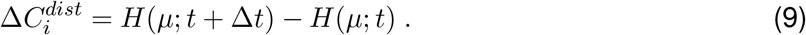

Higher memory noise leads to lower maintenance cost but increased distortion (Fig. 2B). The optimal memory loss from time *t* to *t* + Δ*t* minimizes the weighted sum of distortion cost and maintenance cost,

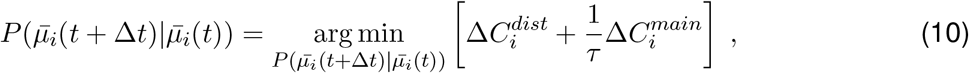

where *τ* is a time constant that controls the relative weight of effort to performance. It has the unit of time to normalize 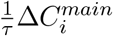 to have the unit of information. Larger values of 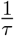 make memory maintenance more costly, resulting in higher distortion; and vice versa. *P*_*i*_(*µ*) undergoes this iterative process from start *t* = 0 to end *t* = *T*_*i*_ of the inter-stimulus interval, resulting in 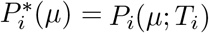. This formalism is general and numerically solvable for arbitrary generative, encoding and maintenance noise models. For the chosen Gaussian sampling noise of our experiment, and the additional assumption of Gaussian noise in encoding and memory drift, we can derive analytical solutions for the optimization (see Methods).

At the end of the trial, the observer makes an estimate 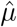 of the generative mean based on the posterior 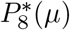. We assume a distance sensitive loss function (ℒ_2_ norm) because in the experiment, subjects received feedback in form of their estimation error in every trial. Therefore, the optimal estimate is the mean of the posterior:

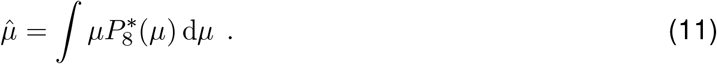

For simplicity, we do not consider motor noise.

### Model behavior

The model explains many different aspects of our specific behavior results, but also makes predictions beyond. First, it offers a unifying account of the well documented variability in regression weighting profiles observed across individual observers and task conditions, which range from emphasizing early samples (primacy) to late samples (recency) in the evidence sequence^25^. With only two free parameters, our model is tightly constrained yet shows a high degree of flexibility in predicting the full range of profiles. As illustrated in Fig. 3A, the model predicts specific profiles depending on how costly the encoding and memory maintenance processes are for the observer. If encoding is costly (large *α*) then it is beneficial for the observer to encode early samples with relative high precision as they are more effective in reducing the uncertainty in the generative mean compared to late samples (Fig. 3B); this leads to a primacy bias. However, large maintenance costs (large 1*/τ*) lead to faster memory decay (Fig. 3G vs. C) and more information loss (Fig. 3H vs. D), thus negate the information benefits of early samples. Eventually, this trade-off results in recency behavior when it is advantageous to encode late samples with relative high accuracy (Fig. 3F). Thus, according to our model, primacy and recency biases result from individual differences in effort allocation between encoding and memory maintenance. Primacy observers put more effort in retaining information in memory while encoding less new information; recency observers lose more information during memory but put more effort in encoding new information. Therefore, different (*α, τ*) value combinations can lead to the same amount of evidence accumulated at the end of a trial, resulting in identical task performance (see also Fig. 4 G).

**Figure 3:**
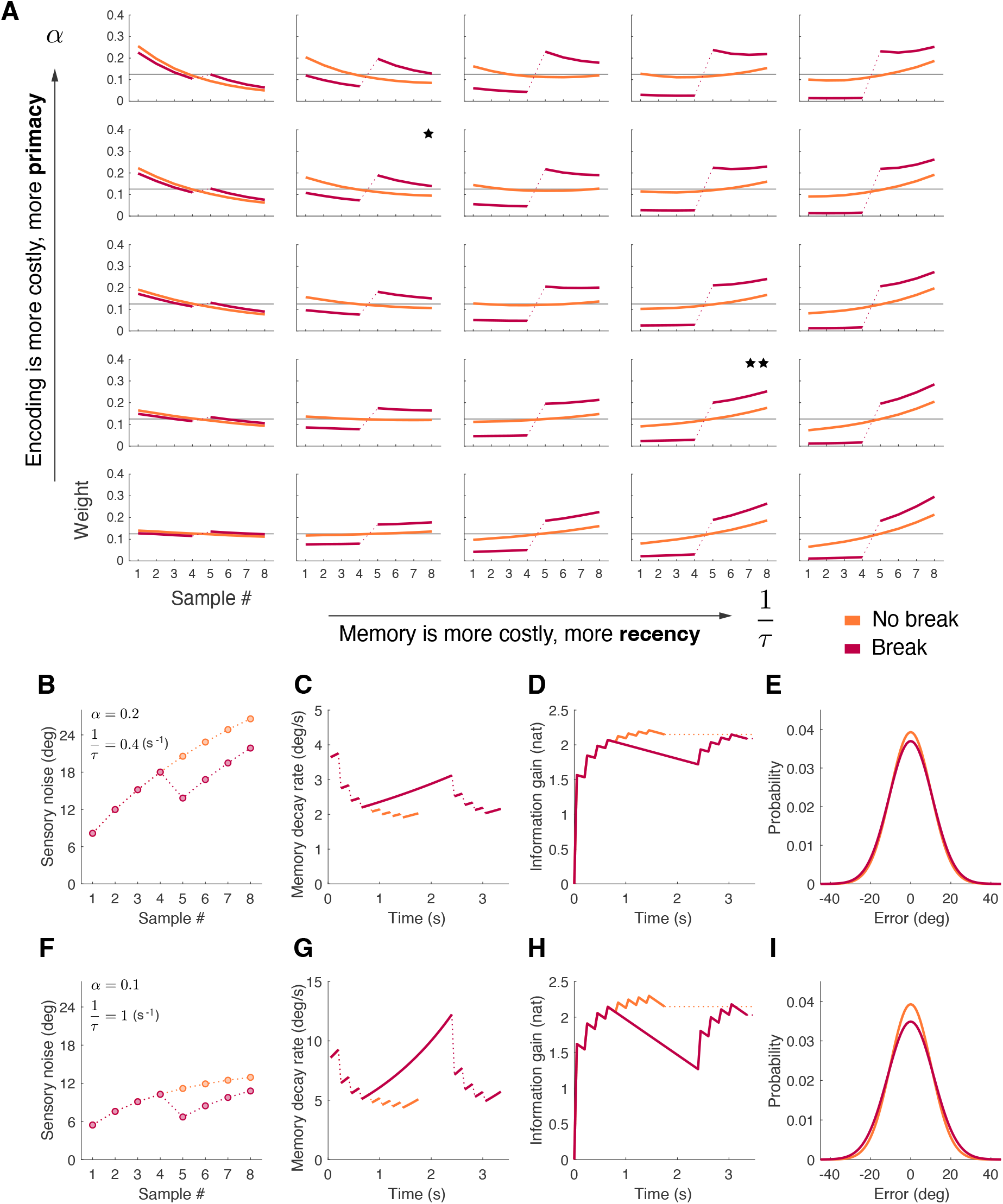
Model behavior (simulations). (A) Predicted regression weights for different value pairs of cost parameters. Increasing values of the memory parameter 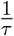 correspond to higher memory maintenance costs and produce recency weighting profiles. Increasing values of the encoding parameter *α* correspond to higher encoding costs and produce primacy weighting profiles. Lower left corner approximates the Ideal observer (*α*, 1*/τ* → 0).The predicted weighting profiles show always positive weight increases for samples after a temporal break in the sample sequence (peak-after-break effect). Panels (B)-(E) show the model dynamics for “Break” and “No break” conditions, assuming a specific parameter pair corresponding to a primacy weighting profile (marked with one star in A). (B) Sensory encoding noise level (SD) with which each sample in the sequence is encoded; Eq.(42). (C) Memory decay rate during the inter-stimulus intervals (and the break); Eq.(52). (D) Information gain/loss throughout the trial; Eq.(53). (E) Predicted estimation error distributions. Panel (F)-(I) show the same for a different parameter value pair that corresponds to a recency profile (marked with two stars in A).

**Figure 4:**
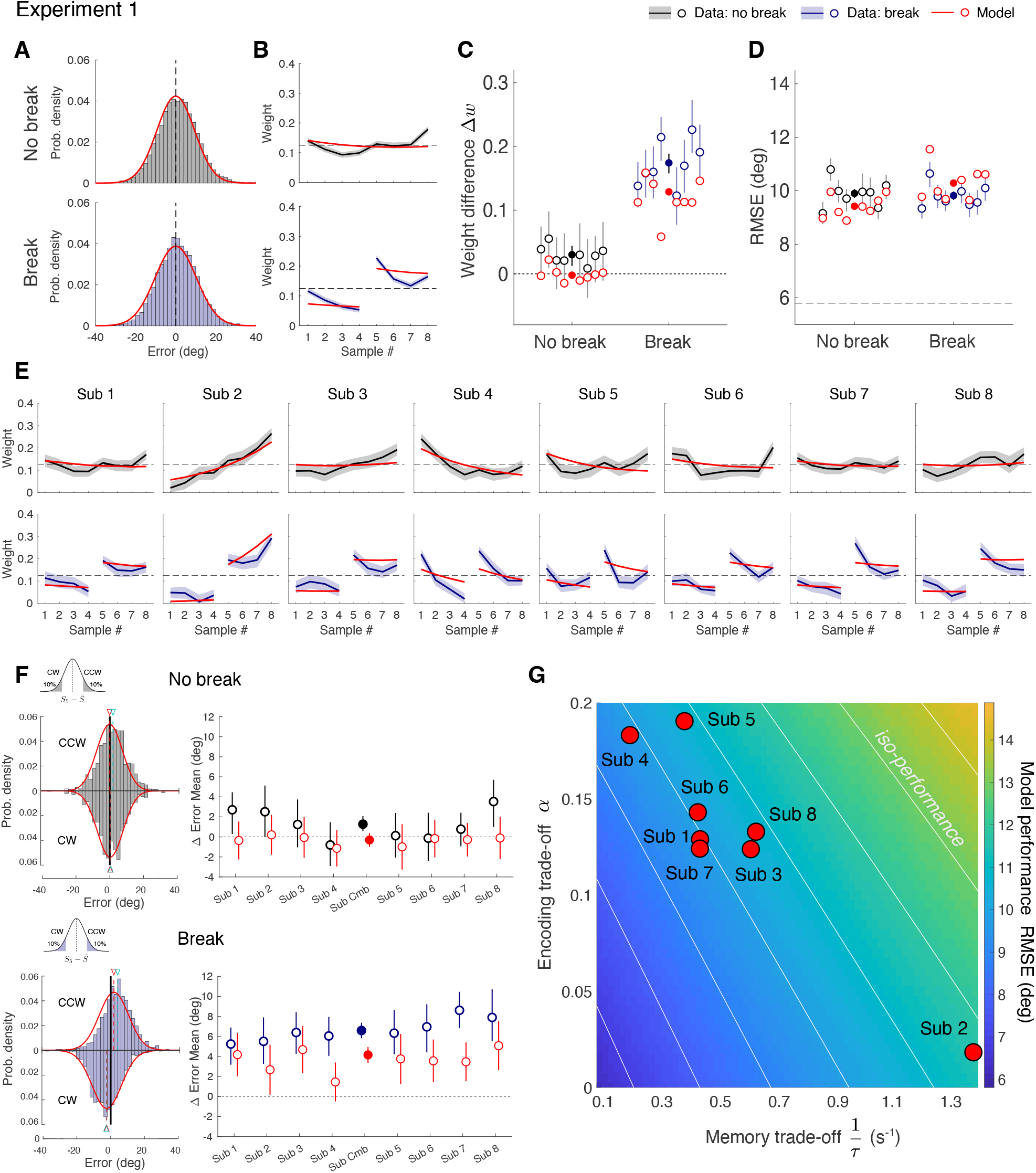
Model fit to data from Experiment 1 (data combined across non-interleaved and inter-leaved blocks). (A) Error distributions for “No break” (black) and “Break” (blue) trials, along with the fit model predictions (red). (B) Circular regression weights and model predictions (combined subject). C) Regression weight differences Δ*w* between the fifth and fourth stimulus sample for all individual subjects (filled circle: combined subject). (D) Estimation performance (RMSE) for “No break” and “Break” trials for individual subjects and their corresponding best fit models (filled circle: combined subject). (E) Regression weights for both conditions; individual subjects. Red curves represent model predicted weights for individual subjects. (F) Left: Error distributions of estimates relative to the sample mean for the two subsets of trials for which the fifth sample was in the 10th percentile tail on either side (CW or CCW); combined subject. Red curves represents model prediction. Cyan and red dashed lines with arrowheads indicate the mean positions of the data and model, respectively. Right: Difference in mean response error between the CCW and CW subsets for each of the eight individual subjects (filled circle: combined subject). Model predictions in red. Note that the subsets across both conditions contain identical trial sequences, and thus are directly comparable. Also, all model predictions are based on a joint fit to all data across both conditions. (G) Subjects’ best fit model parameter pairs. The heatmap represents the model predicted performance (RMSE) for the “Break” condition across the parameter space, with white lines representing parameter manifolds with identical performance levels (Pareto frontiers). Error bars in all panels represent 95% confidence intervals (data: 200 bootstrapped resamples; model: ±1.96 SE of the difference in group means). Supplementary Table S1: Fit model parameters. Supplementary Fig. S1: Comparison between non-interleaved and interleaved trial conditions. Supplementary Fig. S2: Error distributions and model fits for individual subjects.

Second, the model accounts for the observed peak-after-break effect. In order to make up for the information loss during the maintenance of the belief during the break, the model invests more effort to better encode the samples immediately after the break (Fig. 3B, F). This results in higher predicted regression weights for samples after the break. Note that the weight change is always positive, independent of the specific values of the cost parameters, although the weight increase is larger for higher memory costs (larger 1*/τ*).

Finally, the model explains why the estimation performance does not significantly suffer from the information loss during the break (Fig. 3C, G): the loss is almost fully compensated for by the higher encoding precision of the samples after the break such that at the end of the trial the total information gained is approximately identical (Fig. 3D, H). Evidence accumulation is “catching up” on the information gain with the uninterrupted sequence, eventually leading to an almost identical error distribution whose variances represent the RMSEs (Fig. 3E, I).

### Model fits

Our model specifies the probability distribution of the optimal estimate of the generative mean 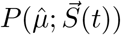 underlying any specific sample sequence 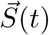 with its individual timing schedule. Given the simplifying assumptions about motor noise and estimation loss (ℒ_2_-norm), this distribution represents the distribution of the posterior mean after the last sample (see Fig. 2C). We can fit the model with its two free model parameters *α* and *τ* by minimizing the negative log-likelihood of the model given the observed data.

We jointly fit the model to data from both the “No break” and “Break” conditions for individual subjects as well as the combined subject (i.e., fit to the combined data from all individual subjects). Based on the fit model parameters, we can analytically specify the corresponding weighting profiles (see Methods). The results shown in Fig. 4 demonstrate that our model successfully accounts for all key characteristics of subjects’ estimation behavior across both conditions. It well captures the response distributions both of the combined subject (Fig. 4A) as well as of individual subjects (Supp Fig. S2A). It also effectively predicts the temporal weighting kernels of the combined (Fig. 4B) as well as of individual subjects with their individually distinct recency and primacy biases (Fig. 4E). It is important to recall that the model was fit to subjects’ estimation reports. The shown weighting kernels are emergent predictions of the fit model. The model further predicts the peak-after-break effect, showing a strong positive weight difference Δ*w* between the samples presented before and after the break (Fig. 4C). Finally, the model also matches subjects’ individual estimation performances, including the finding that there is no significant difference between “Break” and “No break” trials (Fig. 4D).

The peak-after-break effect can be directly observed in subjects’ estimation behavior. To this end, we divided all trials into two subsets based on whether the fifth sample (first post-break sample in the “Break” condition) was clockwise (CW) or counter-clockwise (CCW) of the sample mean of that trial. We limited each set to only contain trials for which the fifth sample was far away from the sample mean, i.e., within the 10th percentile on either side of the distribution. Both subsets contained identical trial sequences across “No break” and “Break” conditions. The estimate distributions for the two subsets are noticeably shifted away from the sample mean in opposite directions in the “Break” compared to the “No break” conditions (Fig. 4F). This shift directly corresponds to the increased regression weight for the first sample after the break (peak-after-break) and is well captured by the model across individual subjects.

### Model predictions for different break durations and positions

A key property of our model is that it can predict accumulation behavior for evidence sequences with arbitrary schedule. We make two general predictions with regard to this schedule (Fig. 5). First, we predict that a longer break duration leads to a larger peak-after-break effect while the total accumulated information, and thus estimation performance, again, remains largely the same (Fig. 5A-C).

**Figure 5:**
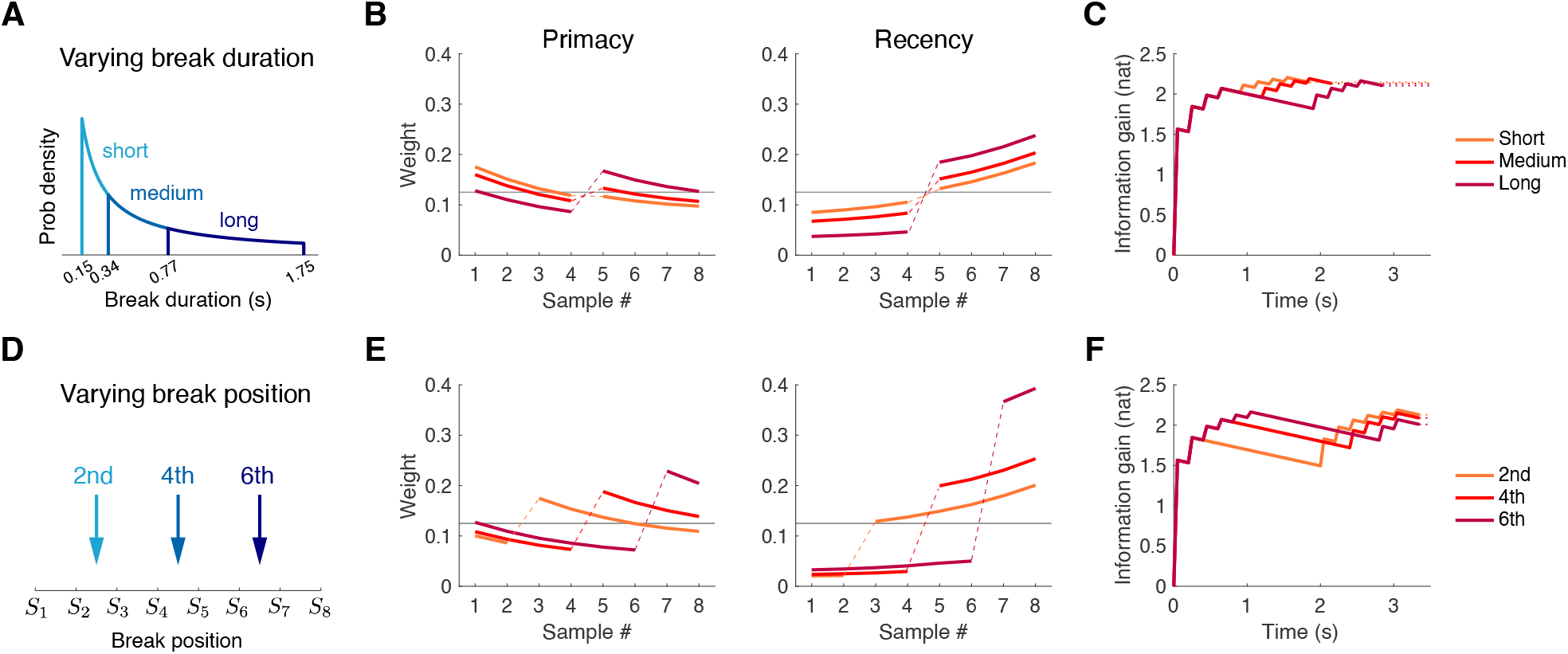
Model predictions for different sequence schedules. (A) Experiment 2: Varying break durations. Break durations were sampled from a logarithmically uniform distribution and categorized into three categories: short (0.15–0.34 s), medium (0.34–0.77 s), and long (0.77–1.75 s) break durations. (B) Predicted weighting profiles for the same primacy and recency example subjects as in Fig. 3. Simulations used break durations of 250 ms, 550 ms, and 1250 ms as representative values for the short, medium, and long categories. (C) Predicted information gain throughout the trial (illustrated for the primacy subject). Total information after the last sample (i.e., estimation performance) is essentially independent on break duration. (D) Experiment 3: Varying break positions. Trials were equally likely to have a break (fixed 1.75 s duration) either after the second, fourth, or sixth stimulus sample. (E) Predicted weighting profiles for the same primacy and recency example subject as in B. (F) Predicted information grain for the example primacy subject. Late breaks are predicted to reduce total information and thus are expected to lead to lower estimation performance (higher RMSE).

Second, we predict that the peak-after-break effect persists regardless of the position of the break in the sequence, yet breaks later in the sequence lead to larger peaks. Also, performance will start to decline because with late breaks there are fewer samples left in the sequence to make up for the memory loss during the break (Fig. 5D-F). We empirically validated both predictions with Experiments 2 and 3.

### Experiment 2: Varying break durations

Experiment 2 was identical to Experiment 1 except that the break after the fourth sample was of variable duration *T* as outline in Fig. 5A. The minimum break duration was set at the regular inter-stimulus interval of 150 ms, and the maximum corresponded to the 1.75 s break duration used in Experiment 1. Because time perception operates on an approximately logarithmic scale^26,27,28,29,30^, we sampled *T* from a log-uniform distribution, dividing it into three, logarithmically equal-sized categories (short, medium, long). We also used the same 550 unique base sample sequences as in Experiment 1, individually generated for each subject. Each unique sample sequence was paired with three break durations selected from the short, medium, and long categories, resulting in a total of 1650 trials per subject. For every trial, break duration *T* was randomly drawn from one of three categories. Eight subjects participated in Experiment 2, each completing 10 testing blocks of 165 trials after training.

Circular regression analysis of the data revealed that the peak-after-break effect occurred independently of break duration (Fig. 6A). However, longer breaks led to larger peaks (Fig. 6B), while estimation performance remained relatively stable across different break durations (Fig. 6C). These results confirm the predictions (Fig. 5). Joint fits of the model across all break durations accurately capture the observed increase in Δ*w* as well as the relative stable estimation performance for different break durations.

**Figure 6:**
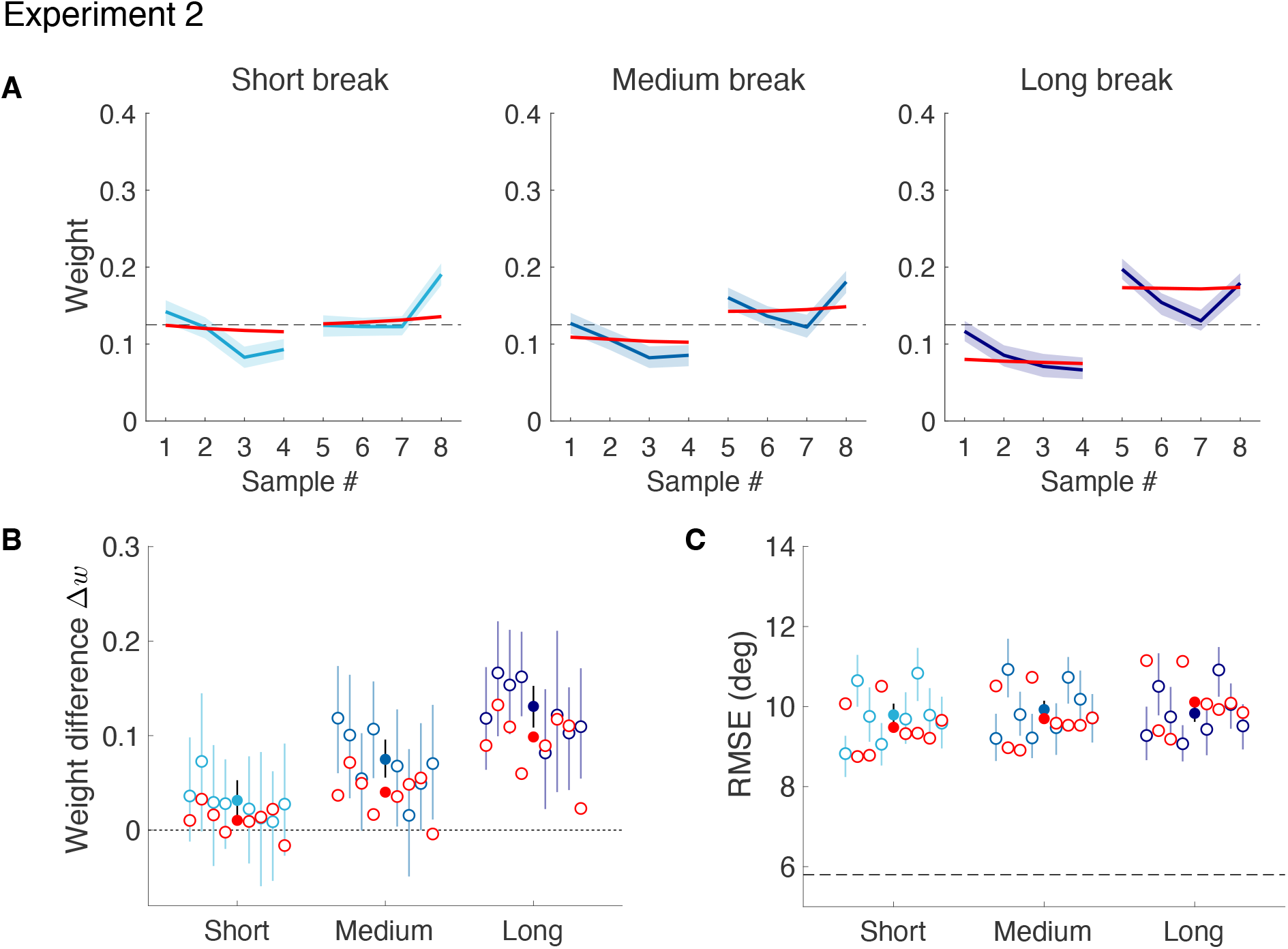
Experiment 2: varying break durations. (A) Regression weights for short, medium, and long break duration trials (combined subject). Red curves show the predicted weighting profiles of the jointly fit model (one parameter pair for all data). (B) Weight differences Δ*w* between the 5th and 4th stimulus sample for individual subjects (filled circles: combined subject), based on behavioral data (black, dark blue) and model fits (red). (C) Estimation performance (RMSE) of individual subjects (filled circles: combined subject) in short, medium, and long break duration trials; behavioral data (black, dark blue) and the joint fit model (red). Dashed line represents the ideal observer performance. Shaded areas and error bars in all panels represent the 95% confidence interval computed over 200 bootstrap samples of the data. The relative positions of data points shown for each duration category in (B) and (C) corresponds to the identity of each subject. Supplementary Tables S1: Fit model parameters. Supplementary Fig. S2B: Error distributions and model fits for individual subjects. Supplementary Fig. S3A: Weighting profiles of individual subjects.

### Experiment 3: Varying break positions

Experiment 3 had the same experimental design as Experiment 1, except that the break position was randomly selected to be either after the second, fourth, or sixth stimulus sample (Fig. 5D). As in Experiment 2, we used a fixed base stimulus set as in Experiment 1, pairing each of the 550 unique sequences with the three break positions, yielding 1650 trials per subject. The training and testing phases were identical to those in Experiment 2.

We observed a peak-after-break effect for all break positions (Fig. 7A), with later breaks resulting in larger weight differences (Fig. 7B). Estimation performance decreased as the break was inserted later in the evidence sequence (Fig. 7C). Both effects, weight difference and performance decrease, are consistent across all subjects, and were predicted by the model (Fig. 5). The magnitudes of the effects are subject specific, depending on the integration characteristics of each subject. For example, Subject 2 shows a relatively large increase in weight difference and performance drop with later break position (2nd dots in panels B and C), which coincides with their distinct recency bias (see Supplementary Fig. S3B). In contrast, Subject 5 shows a moderate increase in both, but is also exhibiting a distinct primacy bias. Our model quantitatively accounts for these correlational patterns and also provides a causal interpretation: Recency bias reflects an observer for which memory maintenance is more costly than sample encoding (Fig. 3). On one hand, such observer will necessarily incur larger information loss during the break and more overall loss with later breaks compared to an observer with primacy bias (hence the increased drop in performance). On the other hand, because encoding is cheaper, the observer can better compensate for the loss with more precise encoding of the samples after the break (hence the increase in weight difference).

**Figure 7:**
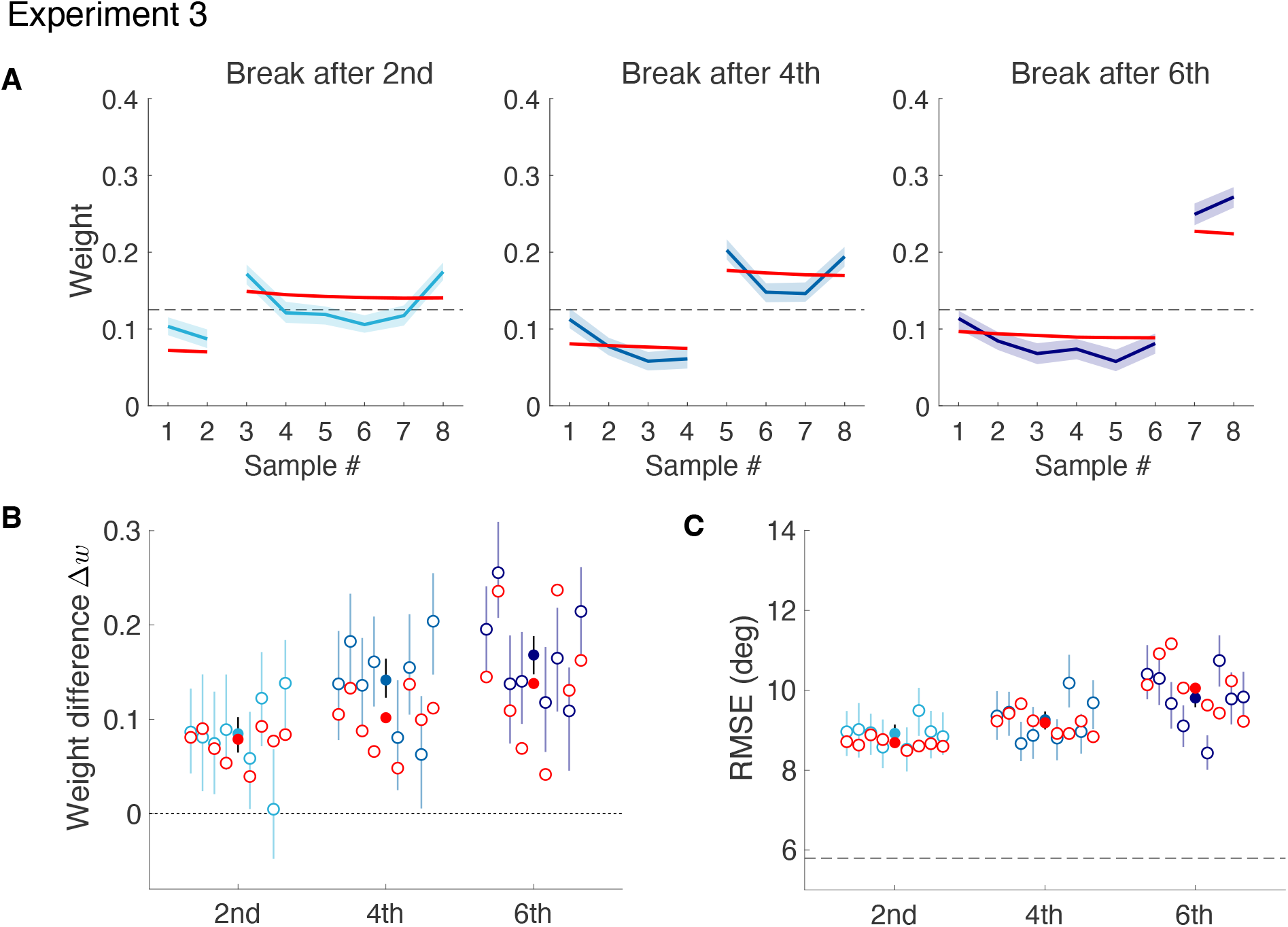
Experiment 3: varying break positions. (A) Circular regression weights for trials with breaks after the second, fourth or sixth stimulus sample. Red curves show the regression weights predicted by the model (combined subject); see Methods. (B) Weight difference Δ*w* between stimuli after and before the break, shown for individual subjects (blue) and the model predictions (red); solid circles represent the combined subject. (C) Estimation performance (RMSE) shown for individual subjects (combined subject, filled circles); data (blue) and individual model predictions (red). Dashed line represents the ideal observer performance (i.e., the sample mean). All above shown model predictions are based on joint fits to data from all conditions. Shaded areas and error bars in all panels represent the 95% confidence interval from 200 bootstrap samples of the data. The relative position of each data point in (B) and (C) corresponds to the identity of each individual subject. Supplementary Tables S1: Fit model parameters. Supplementary Fig. S2C: Error distributions and model fits for individual subjects. Supplementary Fig. S3B: Weighting profiles of individual subjects.

With two free parameters, our model provides a parsimonious account for individual differences in subjects’ behavior across all three presented perturbation experiments based on the single but fundamental assumption that performance-effort trade-offs dynamically control the key processes of evidence accumulation.

### Model ablation analysis

A model ablation analysis illustrates how the PET modules for encoding and memory maintenance each are responsible for different aspects of the integration behavior. Aside from the full model (PET-PET), we considered three reduced model versions. We simulated every version for a few different combinations of parameters and compared their predicted regression weight profiles for the “Break” and “No break” conditions of Experiment 1 (Fig. 8A).

**Figure 8:**
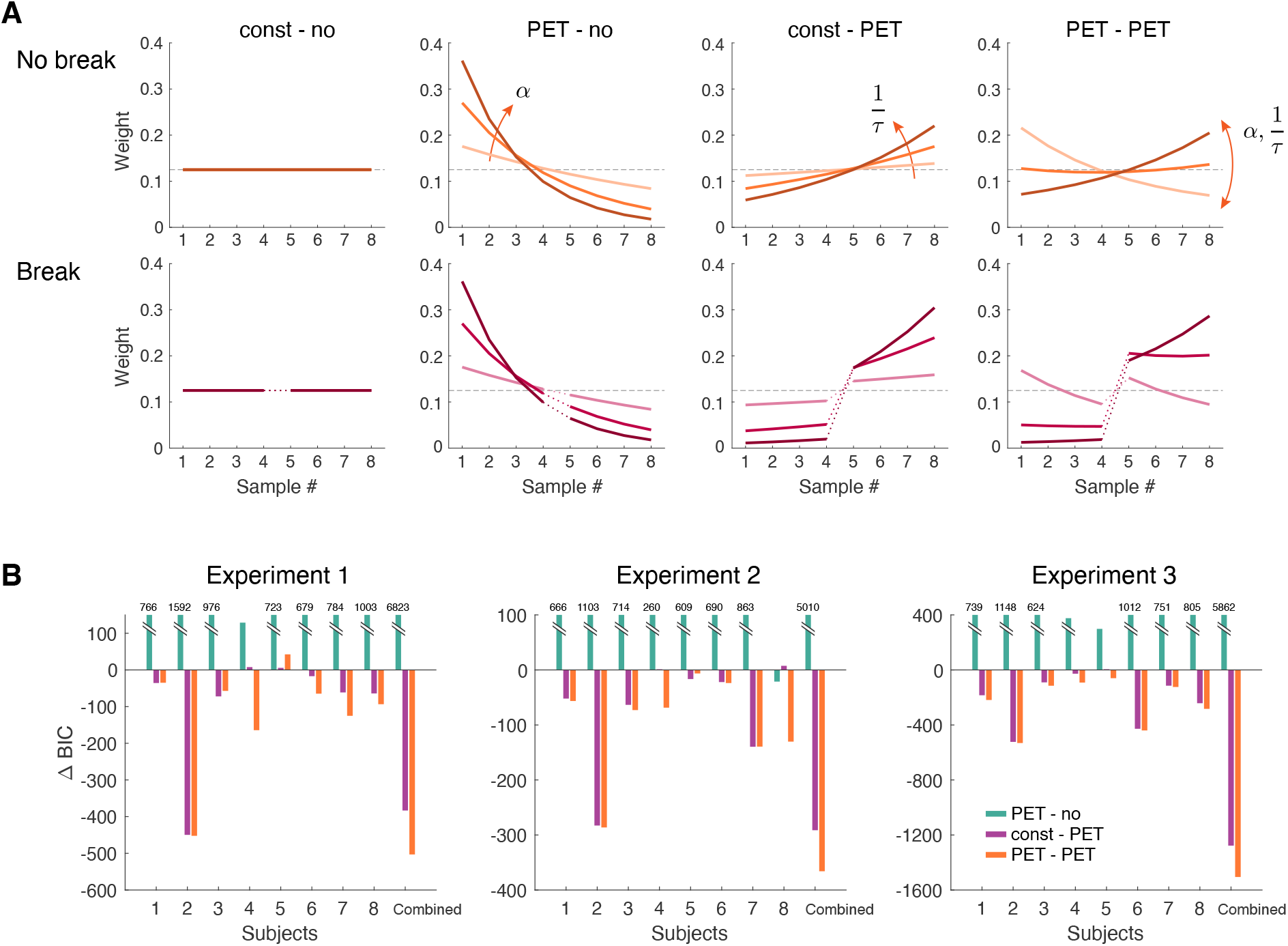
Model ablation analysis. (A) Temporal weighting kernels for the full model (PET-PET) and three ablated versions: the ideal observer model with constant encoding and no memory loss (const-no), a version with only PET controlled encoding but no memory loss (PET-no), and a version with constant encoding and PET controlled memory maintenance (const-PET). Each panel shows the predicted weighting kernels from simulations with different model parameter values in both the “No break” and “Break” conditions. (B) Model fitting ΔBIC values for all best-fit model versions relative to the ideal Bayesian accumulation model (const-no) for individual subjects and the combined subject. Models were jointly fit to data from all conditions within each experiment.

The first model version considered is the Bayesian ideal observer with constant encoding precision and no memory leak (const-no). This model predicts uniform regression weights because every sample is encoded with the same accuracy and maintained without memory loss. A break in the evidence sequence has no effect on the weights. Replacing constant encoding with a PET controlled encoding module results in a model version (PET-no) with lower encoding accuracy for later samples: because the performance gain, i.e., the reduction of uncertainty about the generative mean, decreases for later samples for the same amount of encoding cost, it is not worth encoding them with the same precision as early samples. This consistently leads to primacy behavior for all parameter values (*α*). The weights, however, are identical between “Break” and “No break” conditions because there is no memory loss. In contrast, adding a PET controlled, memory maintenance module to the ideal observer model (const-PET) instead always leads to a loss of information about the earlier samples by the end of the sequence, resulting in a consistent recency bias for all parameter values (1*/τ*). During the duration of the break, the loss accumulates and thus further emphasizes the weights of the samples after the break. This explains, partially, the peak-after-break effect. Only the full model (PET-PET) is able to predict both primacy and recency behavior depending on the specific combination of cost parameter values *α* and 1*/τ* while also consistently predicting the entire peak-after-break effect.

We also tested how well the different model versions can explain our experimental data. We fit all versions to individual subject data from all three psychophysical experiments. The fits were joint fits to the data across all conditions within each experiment. Fig. 8B illustrates the goodness-of-fit comparison between the model versions, measured as the difference in BIC criterion relative to the best fit ideal Bayesian observer model (const-no). Across all three experiments, almost all the subjects are best explained by the full model. Note that although the model version that combines constant encoding with PET controlled memory maintenance (const-PET) is sufficient to account for the data of some recency subjects under some conditions (e.g., Sub 2 in Exp. 1), it fails to explain the data of subjects with primacy behavior (e.g., Sub 4).

## Discussion

With this study, we uncovered previously unknown dynamic aspects of sequential evidence accumulation. We systematically characterized how temporal perturbations (i.e., a break) in otherwise temporally regular evidence sequences affect the accumulation process. We found that evidence accumulation is a much more dynamically controlled, non-stationary process than previously assumed. Subjects’ estimation accuracy remained largely independent of the duration of the break, even though longer break durations should have induced larger memory loss and thus resulted in a performance decrease. Regression analysis revealed a substantial weight increase of the first evidence sample after the break (peak-after-break effect). The magnitude of this effect increased with longer breaks and later break positions, but was consistent across all subjects.

We presented a novel model that provides a rational and intuitive account for all these findings. The model is part of a larger class of resource-rational observer models that are built on the notion that the brain optimally balances performance against cognitive effort^31,32,33,34,35^. Our model provides clearly formulated computational descriptions of all three key aspects of evidence accumulation: encoding, updating, and memory maintenance. Belief updating is Bayesian yet evidence encoding and memory maintenance of the updated beliefs are both dynamically controlled by independent performance-effort trade-offs (PET). These PET modules guarantee that at any moment in time, sensory evidence is only encoded and maintained with the precision required to achieve a level of belief accuracy worth the effort. After a break in the evidence stream, these dynamic control mechanism predict active compensation of the information loss sustained during the break by an enhanced encoding precision for the first sample after the break, which explains the peak-after-break effect and the fact that subjects’ estimation accuracy remained independent on the duration of the break.

Embedding resource constraints (e.g., efficient coding) within the optimal Bayesian observer framework has previously been quite successful in providing intuitive and rational interpretations of otherwise difficult to explain (perceptual) behavior^36,37,38,39^. The current model is extending this general approach to the temporal domain. It also introduces a formulation that dynamically balances optimal resource allocation. Other studies have attributed apparent suboptimality in evidence accumulation to imprecision in Bayesian inference^13,15,40^. In the context of our study, incorporating inference noise would simply add uncertainty whose effect on the posterior belief would be indistinguishable from additional encoding or memory noise; hence its contribution could not be determined unambiguously. By limiting the effects of resource constraints to the representational level, we can maintain an optimal, normative account and interpretation of human behavior. With two free parameters, our model demonstrates a parsimonious yet time-continuous description of individual subjects accumulation behavior across all our experiments. It also expresses subjects’ individual tendencies for primacy or recency bias as their individual efficiency difference between evidence encoding and memory maintenance. Our model is general given its information-theoretic formulation, and thus should be directly applicable to other tasks involving evidence encoding and maintenance.

### Related work

We are unaware of prior work that systematically evaluated the effects of temporal perturbations on evidence accumulation. A recent study has investigated the effect of a single fixed temporal information gap on integration behavior, although in a binary perceptual decision task^41^. The results indicate that subjects overemphasized the information provided by the evidence immediately after the gap, which aligns with our findings and suggests that the effect is general and independent of the specific task.

Several models have proposed computational accounts for recency and primacy biases in evidence accumulation. For example, the approximate inference model^15^ assumes that primacy is the byproduct of a form of confirmation bias induced by a categorical top-down prior that itself is continually updated by previous evidence (essentially counting past information twice); a leaky memory stage then is assumed to partly counteract this bias and shift the profile toward a flat or a recency profile. Drift-diffusion models and their leaky variants instead link different weighting profiles to different types of decision bounds^42,43,44,12,23^. All of these models address binary (i.e., discrete) decision tasks and are not directly applicable to the continuous estimation task of our study. Attractor network models have also been suggested as biophysical equivalents of the accumulation dynamics^45,46,47^. Specifically, the bump attractor network^16^ was recently proposed to model evidence accumulation for a continuous latent state variable. We simulated the bump attractor model for the trial sequences of Experiment 1 for both “break” and “no break” conditions (see Supplementary Fig. S4). Although the network model can mimic integration dynamics that reflect primacy or recency behavior depending on its configuration parameters, it can not account for any of the other characteristic findings of our perturbation experiments. In particular, rather than exhibiting a consistent positive peak-after-break effect it predicts a weight change that can be positive or negative depending on the overall integration characteristic. Furthermore, it predicts a substantial performance drop for the “break” condition rather than the maintained performance level seen in our data. These findings demonstrate that a model with stationary dynamics cannot account for our behavioral data.

Two recently proposed resource-rational models propose a similar trade-off formulation for Bayesian belief updating as our model albeit again for a discrete decision variables^48,49^. However, both models assume that resource limitations lead to imperfect Bayesian inference instead of less accurate encoding. Also, they focus on belief updating only, and ignore the costs associated with maintaining beliefs over time. As a result, these models are not defined on continuous time and thus can not account for evidence accumulation of sequences with irregular schedule. In contrast, another recent study proposed a trade-off model for optimally controlling the imprecision of memory maintenance of evidence similar to our model, yet assumed evidence accumulation to maintain fixed encoding precision^50^. The results of this study illustrate that PET controlled memory maintenance can explain interesting recency effects in economic forecasting. However, as the model ablation analysis has shown, both PET controlled encoding and memory maintenance are necessary to account for recency and primacy behavior, and more generally, to provide a complete and accurate explanation of the presented experimental data (Fig. 8).

### Outlook

Our results give rise to potential future research directions. First, it is worth noting that our general experimental design would have profited from a mask stimulus after the final sample. Because every other sample was effectively masked by the next sample in the sequence, the lack of a mask has likely given the final sample an encoding advantage^51^. Indeed, the results of our regression analysis indicate that most subjects assigned a disproportionately high weight to the final sample (e.g., Fig. 4E) which is also very consistently present for the combined subject data (Figs. 4B, 6A and 7A). The model would have likely even better explained the data with a mask. Thus future experiments of similar design should include a mask after the final evidence sample.

Second, while we tested evidence accumulation for a continuous latent variable, many perceptual tasks involve inference over a hierarchical generative model with higher-level categorical representations. It will be interesting to extend our model to such hierarchical representations, and explore to what degree observed and seemingly irrational interactions between the representational levels, such as confirmation biases^52,53^ or holistic perception^54^, can be explained as the result of optimal performance-effort trade-offs.

Gaining a better understanding of the aspects that determine the trade-off parameter values of the PET modules for individual subjects is another potential future direction. We have shown that subjects have different parameter values, reflecting their individual evidence accumulation preferences and efficiencies. What factors could lead to changes of these values, and how? Recent results have suggested that the level of complexity with which human perform cognitive tasks is correlated with the overall metabolic resources available, with simpler strategies chosen at lower resource levels^55,56^. We speculate that manipulations of the physiological energy state can lead to systematic changes in subjects’ performance-effort balances, reflected in changes to the trade-off parameter values of our model. State dependent differences in the efficiency of sensory encoding and memory maintenance may also explain the observed differences in sensory integration behavior^57^ and decision-making difficulties^58,59,60^ between clinical (e.g., autism spectrum conditions, schizophrenia) and control groups. This hypothesis is qualitatively supported by reported alterations in visual stimulus encoding capacity^61,62,63^ and impaired working memory^64,65^ of clinical groups. Trade-off balances may also adapt to the overall information content provided by the evidence sequence. Changing the sampling variance in our experiments (see Fig. 1A), for example, reduces the information content of each sample, which then automatically leads to a lower encoding precision based on our PET encoding process (Eqs. (3)–(8)). However, the system as a whole may try to compensate for the overall lower information content by being prepared to invest more effort (lower *α*). Alternatively, the system may be even less willing to invest effort as the low information content is deemed not worthy (hence, higher *α*). We are currently running behavioral experiments to distinguish between these two hypotheses.

Finally, it will be interesting to link our model formulation to physiological measures and signals. Our model is formulated at an information-theoretic level. This has the advantage that it provides a general, implementation-independent computational solution to the optimization problem. It is mathematically tractable and, for the Gaussian case, even allows for closed-form solutions. Mutual information is also a quantity that we can extract from measurements of neural population activity^53^. However, studies will be necessary to determine how well the model’s information based effort metrics compare to actual metabolic costs. Irrespective, the model provides detailed predictions of the trajectories of many biophysical (e.g., sensory and memory noise) and decision relevant variables (e.g., posterior beliefs) throughout individual trials. These time-continuous trajectories provide rich and fine-grained dynamic model predictions that will help to identify the equivalent neural processes underlying evidence accumulation based on large scale neuroimaging measurements of neural activity during evidence accumulation^66^.

## Acknowledgments

This work emerged from a collaborative project (CRCNS) that was supported by the National Science Foundation (NSF), project IIS-1912232 and the German Federal Ministry of Education and Research (BMBF) project 01GQ1907. Partial support also came from the NSF and the Department of Defense (DoD) OUSD (R&E) under cooperative agreement PHY-2229929 (The NSF AI Institute for Artificial and Natural Intelligence ARNI). Preliminary results of the work have been presented a the Vision Science Society Annual Meetings^67,68^. We thank the members of the Computational Perception and Cognition Laboratory for the many insightful discussions.

## Methods

### Psychophysical experiments

Fourteen subjects with normal or corrected-to-normal vision (eight male, six female; two non-naïve; age range 21–39, mean: 27.8 ± 5.2 years) participated in the experiments. Five subjects (Sub 1-5, one male, four female; two non-naïve) completed all three experiments, while the remaining subjects took part in only one of the three experiments. All subjects provided informed consent. The experiments were approved by the Institutional Review Board of the University of Pennsylvania under protocol #833347.

#### General methods

Subjects were seated in a dimmed room in front of a specialized computer monitor (VIEWPixx3D) with a 120 Hz refresh rate and a resolution of 1920 × 1080 pixels. The viewing distance from the screen to the eyes was maintained at 85 cm using a chin rest. All experiments were programmed in MATLAB (MathWorks. Inc.) using Psychtoolbox-3 (http://psychtoolbox.org^69^) for stimulus generation and presentation. The code was run on an Apple Mac Pro computer with Quad-Core Intel Xeon 2.93 GHz, 8 GB RAM.

Subjects performed a sequential estimation task as outlined in Fig. 1A. In each trial, subjects first fixated on a black fixation point (diameter: 0.21^°^) at the center of the screen for 50-100 ms. The fixation point then turned blue, and 8 stimulus samples were displayed as white solid dots (diameter: 1^°^) in rapid succession around the central fixation point at 5^°^ visual eccentricity. The angular position of each sample *S*_*i*_ was generated from a Gaussian distribution with a trial-specific mean *µ*_*j*_ and a fixed variance 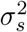:

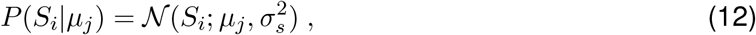

where *i* represents the sample position in the sequence, and *j* indicates the trial number within a block. The fixed variance *σ*_*s*_ was set to 16.3^°^, and the generative mean *µ*_*j*_ was randomly sampled as integers from the full circle.

Each stimulus sample was presented for 50 ms and the regular inter-stimulus interval (ISI) was set to 150 ms, resulting in a 200 ms stimulus onset asynchrony (SOA). Subjects were required to maintain fixation throughout the entire stimulus presentation, signaled by a blue fixation point. Fixation was monitored for the subjects’ dominant eye using an EyeLink 1000 Plus eye tracker.

After stimulus presentation, the fixation point turned black and a 300 ms blank digestion period followed, allowing subjects to consolidate the sequence. Subjects were then asked to provide an estimate of the generative mean of the stimuli distribution by moving a cursor, shown as a red estimation line, using an analog joystick (Sony PS4 Dualshock gamepad). To avoid any initial orientation biases, the cursor only became visible after subjects started moving the joystick. Subjects were allowed up to 10 s to adjust their estimates. Once they finalized their estimate, subjects confirmed it by pressing a button on the gamepad. A confirmation beep (800 Hz, 0.1 s) was played and the true generative mean was displayed as a green line alongside the confirmed red estimation line (see Fig. 1A). Trials were aborted and moved to the end of the block if subjects broke fixation or failed to confirm their estimate within the allotted time. Trials were initiated by a button press.

Before starting the main experiments, subjects performed 14 training trials to familiarize themselves with the visual estimation task. They then proceeded to the formal testing, with every subject completing 2200 trials across 8 blocks for Experiment 1 or 1650 trials across 10 blocks for Experiment 2 and 3. Each block lasted approximately 25 min. The base payment was set to $10 per hour. Subjects were incentivized by the opportunity to earn a monetary bonus based on their overall performance compared to their subject peers ($50, $25, and $10 for best, second and third best performance, respectively).

### Experiment 1: No break vs. Break

Eight subjects participated in Experiment 1. They performed the task under three sessions. In the first session, stimulus samples were presented sequentially with a fixed ISI of 150 ms in each trial (“No break” trials). In the second session, the sequences were interrupted for 1.75 s after the 4th sample (“Break” trials). For each subject, we generated a base stimulus set consisting of 550 unique random sample sequences. In each trial, a sample sequence from the base set was rotated by a random reference angle, drawn uniformly from the full circle. When we refer to “identical stimulus sequences” in the main text, we are referring to these base sequences prior to rotation. Subjects completed 550 “No break” trials in the first session, divided into two blocks of 275 trials. They then performed 550 “Break” trials in the second session, also split into two blocks of 275 trials. In both sessions, trial sequences were generated by shuffling the sample order. The same procedure was used in the third session, in which subjects completed 1100 trials with “No break” and “Break” trials randomly interleaved across four blocks. Thus, each subject was tested on the same 550 base sequences across all conditions and sessions. Before starting the first block of each session, subjects performed 14 training trials.

### Experiment 2: Varying break duration

Eight subjects participated in Experiment 2. Similar to the “Break” trials in Experiment 1, a temporal break was introduced in the middle of an otherwise regular stimulus sequence. The duration of the break, however, varied in the range from 150 ms (regular ISI) to 1.75 s (the break length used in Experiment 1). Specifically, we categorized break durations into three logarithmically uniform distributions. The short break distribution ranged from 150 ms to 340 ms, the medium break distribution from 340 ms to 772 ms, and the long break distribution from 772 ms to 1.75 s. Subjects first completed 14 training trials with break durations randomly drawn from all three distributions. Following training, they completed 10 blocks of 165 trials each, for a total of 1650 trials. Of these, 550 trials were assigned to each break duration distribution. For each subject, trial sequences were generated from the same set of 550 unique base sequences, as described in Experiment 1. In each trial, a base sequence was rotated by adding a randomly sampled reference value.

### Experiment 3: Varying break position

Eight subjects participated in Experiment 3. The procedure was identical to Experiment 2, except that the break duration was fixed at 1.75 s (the same as in the “Break” trials in Experiment 1), but the position of the break varied. Instead of always placing the break in the middle of the sequence (i.e., the 4th ISI), we inserted the break in either the 2nd, 4th, or 6th interval within the sequence. Subjects first completed 14 training trials and then performed the formal testing of 10 blocks with 165 trials each. Trials were evenly divided across the three break positions, with 550 trials assigned to each. Trial sequences were generated as in Experiment 2, using the same set of 550 base sequences for each subject.

### Circular regression

A linear regression model in circular space was used to estimate the relative contribution of each stimulus sample to the estimate of the generative mean. This regression model assumes that the estimated generative mean 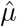 is the angular direction of the sum of vectors whose directions represent the angular position of the samples and whose lengths are the regression weights; thus

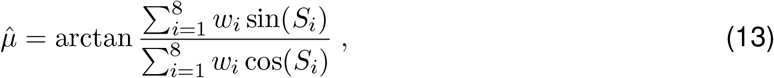

where *w*_*i*_ is the weight of the *i*th stimulus sample *S*_*i*_, with the constraint

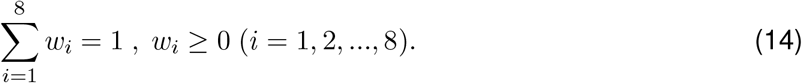

We obtain the maximum likelihood estimate of the weights by minimizing the cosine difference between model prediction and the data:

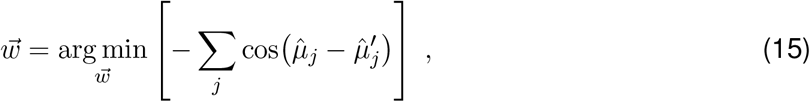

where 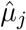 is the model prediction and 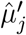 is the subject’s estimate in the *j*th trial.

### Performance Effort Trade-off (PET) model

The model assumes performance effort trade-offs modules for encoding and memory maintenance, respectively. Note that the formulations for both modules are equivalent using mutual information between in- and output as effort, and the reduction in uncertainty in posterior belief as performance metric^24^. In the following, we derive the formulations in detail, and present an analytical solution under the assumption of Gaussian noise.

#### Encoding

The encoding cost (effort; Eq.(3)) is defined as the mutual information between the sensory measurement and the sample:

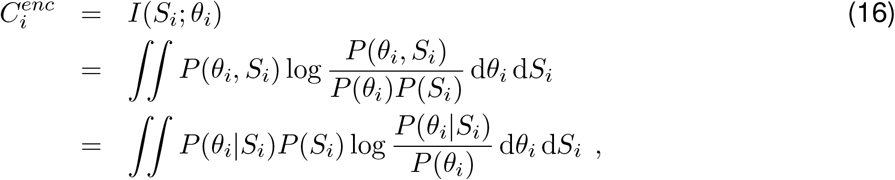

where

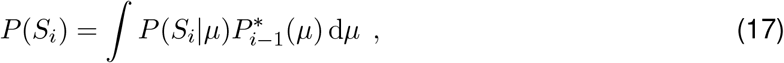

and

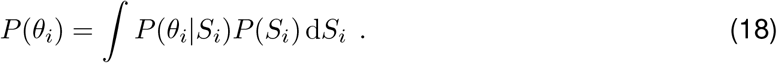

The encoding inaccuracy (performance; Eq.(4)) is defined as the entropy of the generative mean after Bayesian updating:

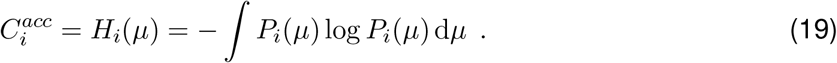

#### Memory maintenance

The incremental memory maintenance cost (Eq.(8)) is defined as the mutual information between the mean of *P*_*i*_(*µ*) before and after the memory loss over a small time step Δ*t*:

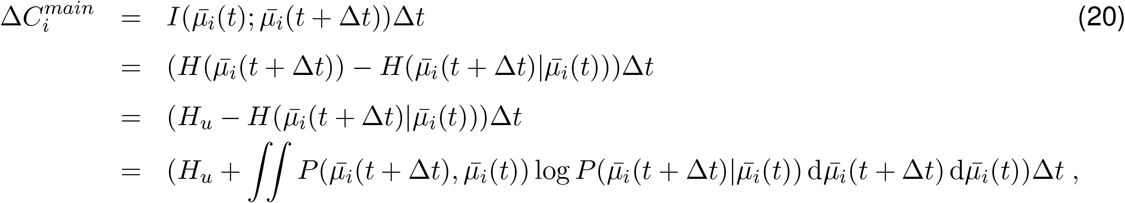

where *H*_*u*_ is the entropy of a uniform distribution. Memory distortion (Eq.(9)) is defined as the increase of entropy of *µ*:

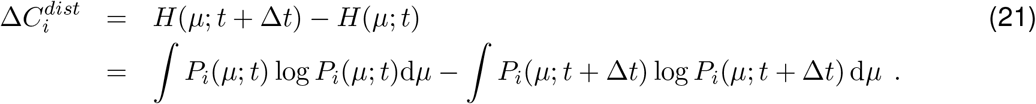

#### Analytical solution under Gaussian noise

Given Gaussian distributed sample sequences in the experiments and the assumption of Gaussian noise in encoding and memory drift, we can derive an analytical solution to the optimization problem. The analytical solution holds under small noise conditions, for which the circular feature space can be approximated by a linear space.

First, we show that the probability distribution of *µ* (belief) remains Gaussian throughout the entire integration process. Assuming Gaussian noise in the sensory measurement

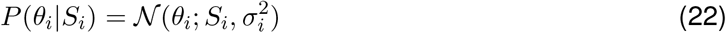

with variance 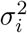, the probability of the measurement *θ*_*i*_ for a given generative mean *µ* is

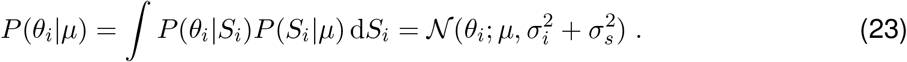

Because *µ* has a uniform prior *P*_0_(*µ*) = 𝒰_[0,2*π*)_, after observing the first sample, the probability of *µ* is updated to a Gaussian distribution:

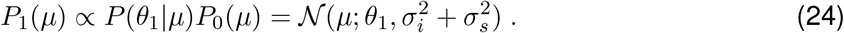

For subsequent samples, if the probability of *µ* before observing the *i*th sample is a Gaussian distribution with width *σ*_*µ*_:

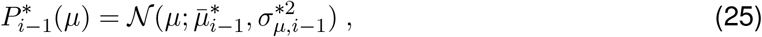

it remains Gaussian after the update with the measurement of the *i*th sample:

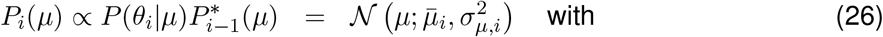

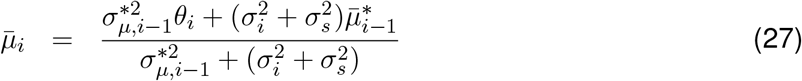

and

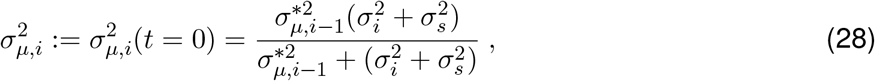

where *t* = 0 represents the start of the memory period (right after evidence updating). During the memory process in the *i*th inter-stimulus interval, we assume the mean of the probability of *µ* drifts according to a Gaussian random walk

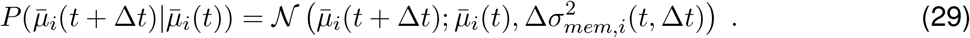

If the probability of *µ* at time *t* is a Gaussian distribution with width *σ*_*µ,i*_(*t*),

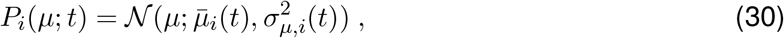

then at time *t* + Δ*t*, it is still Gaussian:

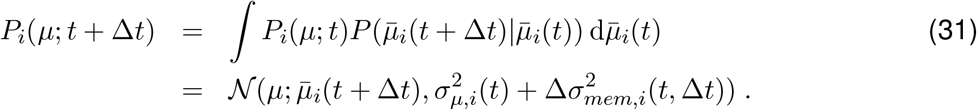

Therefore if the probability of *µ* before the inter-stimulus interval is a Gaussian distribution

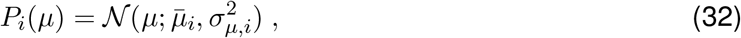

it is still Gaussian after the inter-stimulus interval:

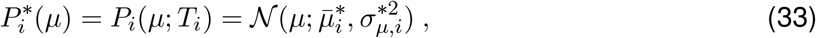

Where

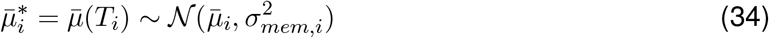

and

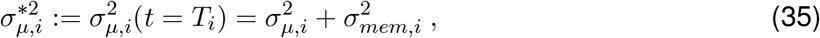

with

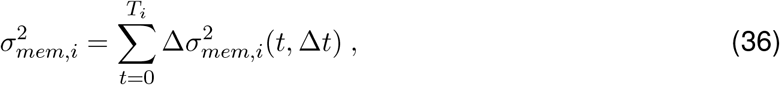

where *T*_*i*_ is the duration of the *i*th inter-stimulus interval. Therefore, the probability of *µ* remains a Gaussian distribution after the encoding of the first sample throughout the rest of the trial.

Next, we derive the encoding and memory noise that optimize the performance effort trade-off. During encoding, with Gaussian encoding noise (Eq.(22)), the encoding cost (Eq.(16)) is

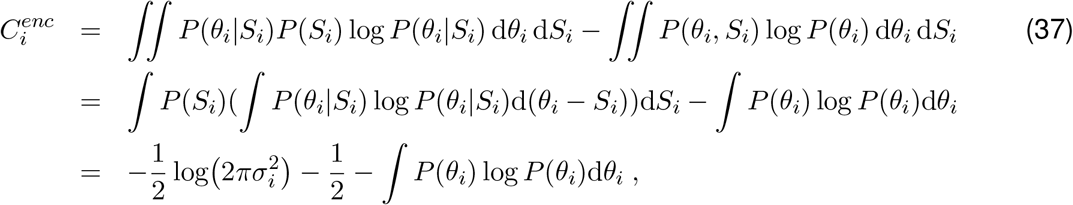

where

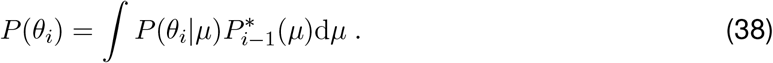

For the first sample, *P*_0_(*µ*) = 𝒰_[0,2*π*)_, so *P*(*θ*_1_) = 𝒰_[0,2*π*)_. For later samples (*i* ≥ 2), following Eq.(23) and Eq.(25),

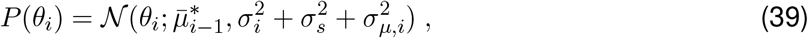

so the encoding cost is

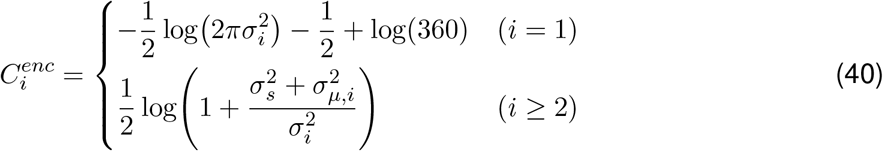

Following the probability of *µ* after encoding the new sample in Eq.(24) and Eq.(26), the inaccuracy (Eq.(19)) is

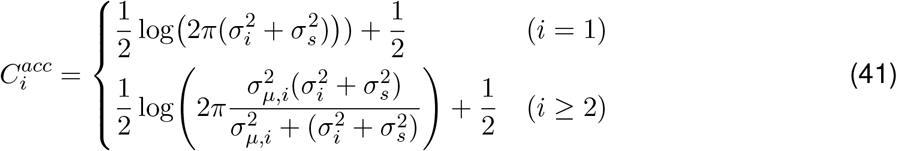

So the optimal encoding noise is

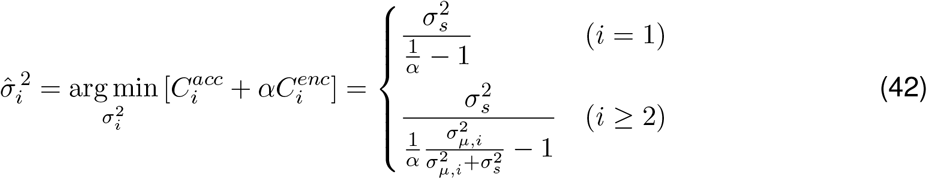

where *σ*_*µ*_ is the standard deviation of 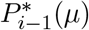 right before encoding the *i*th sample.

During the memory process in the *i*th inter-stimulus interval, based on the assumption of a Gaussian random walk (Eq.(29)-(31)), from *t* to *t* + Δ*t*, the memory distortion cost (Eq.(21)) is

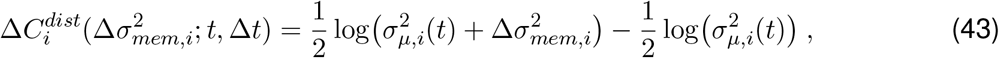

and the memory maintenance cost (Eq.(20)) is

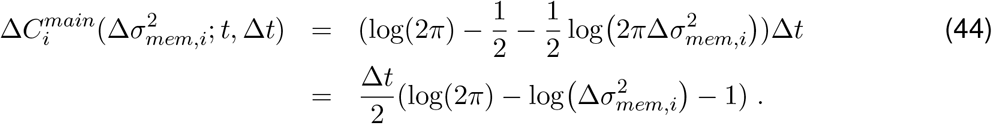

The optimal memory loss is

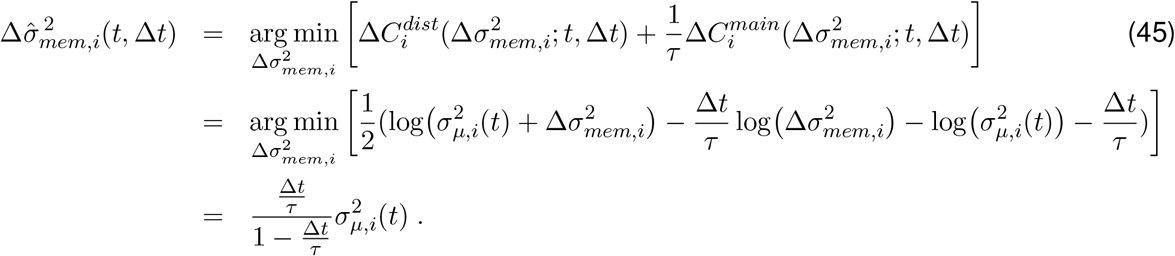

So the width of the probability of *µ* as a function of time can be solved as

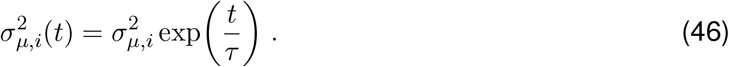

Therefore the probability of *µ* at the end of the memory period is

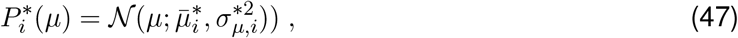

with

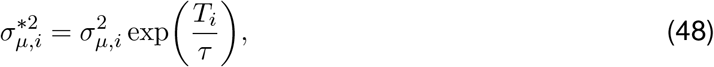

where *T*_*i*_ is the duration of the *i*th inter-stimulus interval, and

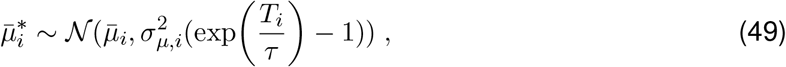

where 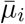 and *σ*_*µ,i*_ are the mean and standard deviation of *P*_*i*_(*µ*) before the memory period.

#### Estimate distribution and weights

For the Gaussian case with analytical solution, the final estimate 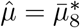 is a linear combination of the sensory measurements of the samples plus memory noise. The probability distribution of the estimate given the true generative mean *µ* is a Gaussian distribution with mean *µ* and the variance of the final posterior 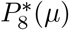,

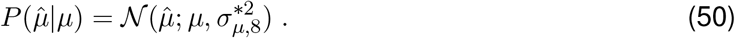

The linear weights of the sensory measurements can be derived iteratively and analytically (Eqs. (24) and (27)). For a given sample sequence 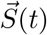, the probability distribution of the estimate is a Gaussian distribution whose mean is the weighted average of the samples, and the variance is 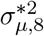 minus the noise induced by stimulus sampling:

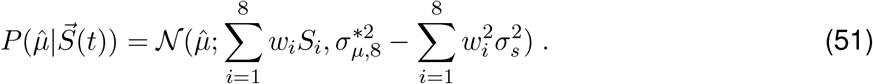

This linear analytical solution holds for circular variables under a small noise condition, which is the case in the present experiment. To facilitate a fair comparison between the circular regression of the experimental data and the model, the weights based on the model fits (Fig. 4A,D; Fig. 6A; Fig. 7A) are circular weights calculated for the experiment trials. The weights of the model predictions (Fig. 3A; Fig. 5B,F) are calculated by simulating 1000 trials.

#### Model behavior

The sensory noise of each sample (Fig. 3B,F) can be calculated through Eq. (42). The memory decay rate (Fig. 3C,G) illustrates the rate of increase of the standard deviation of 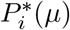. From Eq.(46), the memory decay rate is

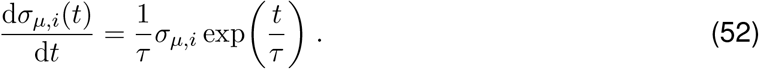

The information gain (Fig. 3D,H; Fig. 5C,F) is the decrease of entropy at every time point compared to the beginning of the trial:

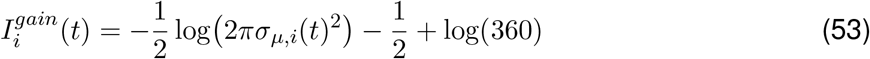

where *σ*_*µ,i*_(*t*)^2^ is the variance of the posterior of *µ* at time *t* in the *i*th inter-stimulus interval. The error distribution (Fig. 3E,I) is a Gaussian distribution centered at 0, with variance 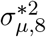 as explained above (Eq.(50)).

## Data and Code availability

All data and code will be made publicly available at the time of publication.

## Supplementary information

### Fit model parameters

**Table S1:** Best-fitting model parameters for individual subjects and the combined subject across all three experiments. For each experiment, parameters were obtained from a joint fit across all conditions within the experiment (e.g., all different break positions in Experiment 3) by minimizing the negative log-likelihood of the model given the data. Parameter *α* specifies the balance between stimulus encoding cost and inaccuracy cost during the encoding phase, while the parameter 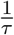 (units: s^−1^) specifies the balance between memory maintenance cost and distortion cost during the memory phase. “Sub Cmb” denotes fits to data combined across all 8 subjects. Note: Sub 1–Sub 5 are subjects who completed all three experiments; all other subjects are unique to a single experiment.

| Experiment 1: No break vs. Break |  |  |  |  |  |  |  |  |  |
| --- | --- | --- | --- | --- | --- | --- | --- | --- | --- |
| Parameter | Sub 1 | Sub 2 | Sub 3 | Sub 4 | Sub 5 | Sub 6 | Sub 7 | Sub 8 | Sub Cmb |
| $\alpha$: encoding | 0.13 | 0.02 | 0.12 | 0.18 | 0.19 | 0.14 | 0.12 | 0.13 | 0.14 |
| $\frac{1}{\tau}$: memory | 0.45 | 1.38 | 0.63 | 0.21 | 0.40 | 0.44 | 0.45 | 0.64 | 0.53 |
| Experiment 2: Varying break duration |  |  |  |  |  |  |  |  |  |
| Parameter | Sub 1 | Sub 2 | Sub 3 | Sub 4 | Sub 5 | Sub 6 | Sub 7 | Sub 8 | Sub Cmb |
| $\alpha$: encoding | 0.09 | 0.04 | 0.10 | 0.13 | 0.13 | 0.17 | 0.06 | 0.21 | 0.13 |
| $\frac{1}{\tau}$: memory | 0.57 | 1.29 | 0.71 | 0.34 | 0.56 | 0.67 | 0.87 | 0.17 | 0.60 |
| Experiment 3: Varying break position |  |  |  |  |  |  |  |  |  |
| Parameter | Sub 1 | Sub 2 | Sub 3 | Sub 4 | Sub 5 | Sub 6 | Sub 7 | Sub 8 | Sub Cmb |
| $\alpha$: encoding | 0.10 | 0.03 | 0.10 | 0.13 | 0.14 | 0.05 | 0.11 | 0.08 | 0.10 |
| $\frac{1}{\tau}$: memory | 0.45 | 0.74 | 0.36 | 0.26 | 0.20 | 0.74 | 0.41 | 0.50 | 0.43 |

**Figure S1:**
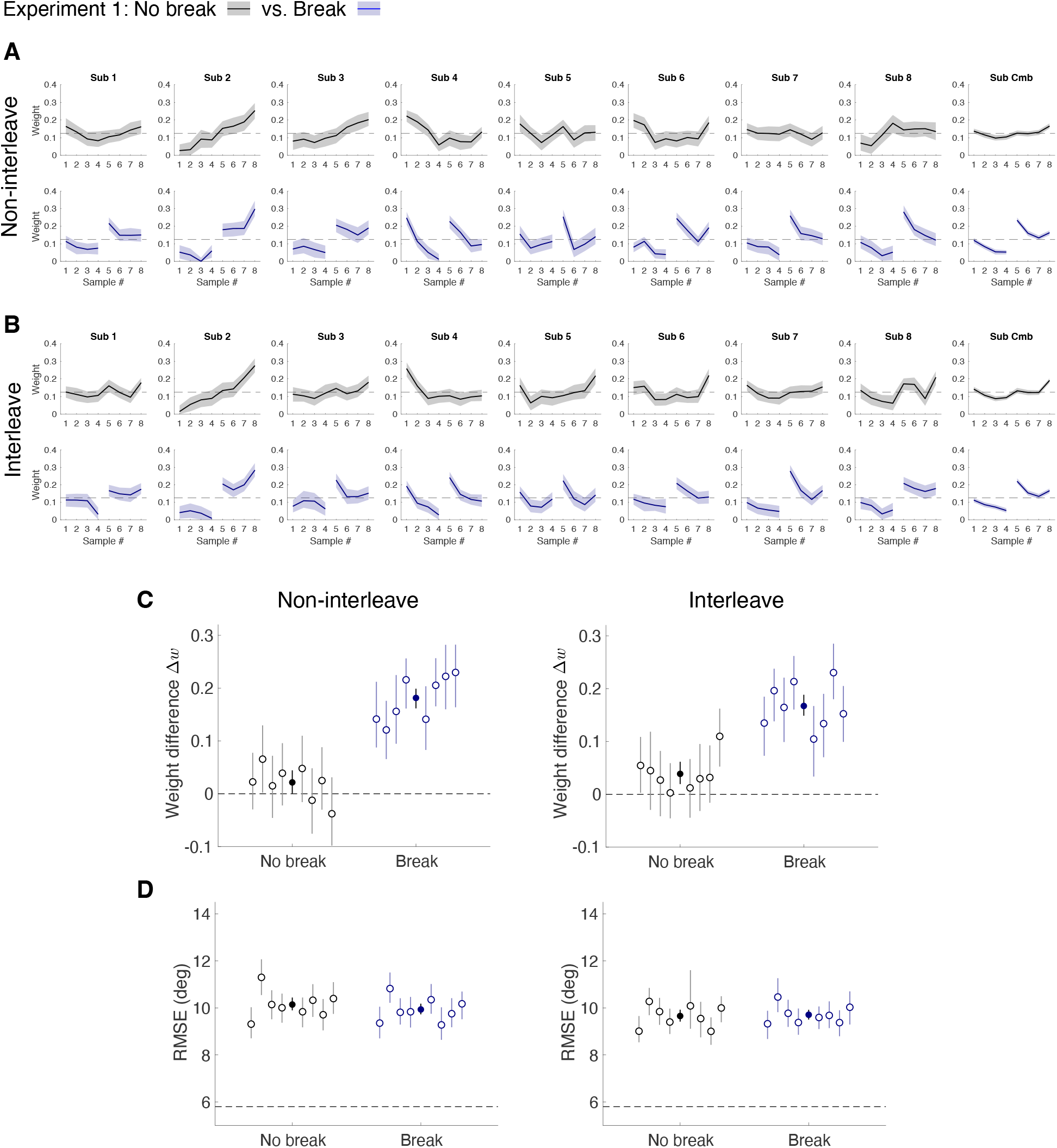
Circular regression weights for “No break” (top row) and “Break” (bottom row) trials in non-interleaved (A) and interleaved blocks (B) of Experiment 1. Shaded areas indicate 95% confidence intervals based on 200 bootstrap results. (C) Weight difference Δ*w* between the 5th and 4th stimulus sample for “No break” and “Break” trials in non-interleaved blocks (left) and interleaved blocks (right). (D) Estimation performance (RMSE) for “No break” and “Break” trials in non-interleaved blocks (left) and interleaved blocks (right). Hollow circles denote individual subjects, and solid circles denote the combined subject. Error bars represent 95% confidence intervals computed from 200 bootstrap samples of the data.

**Figure S2:**
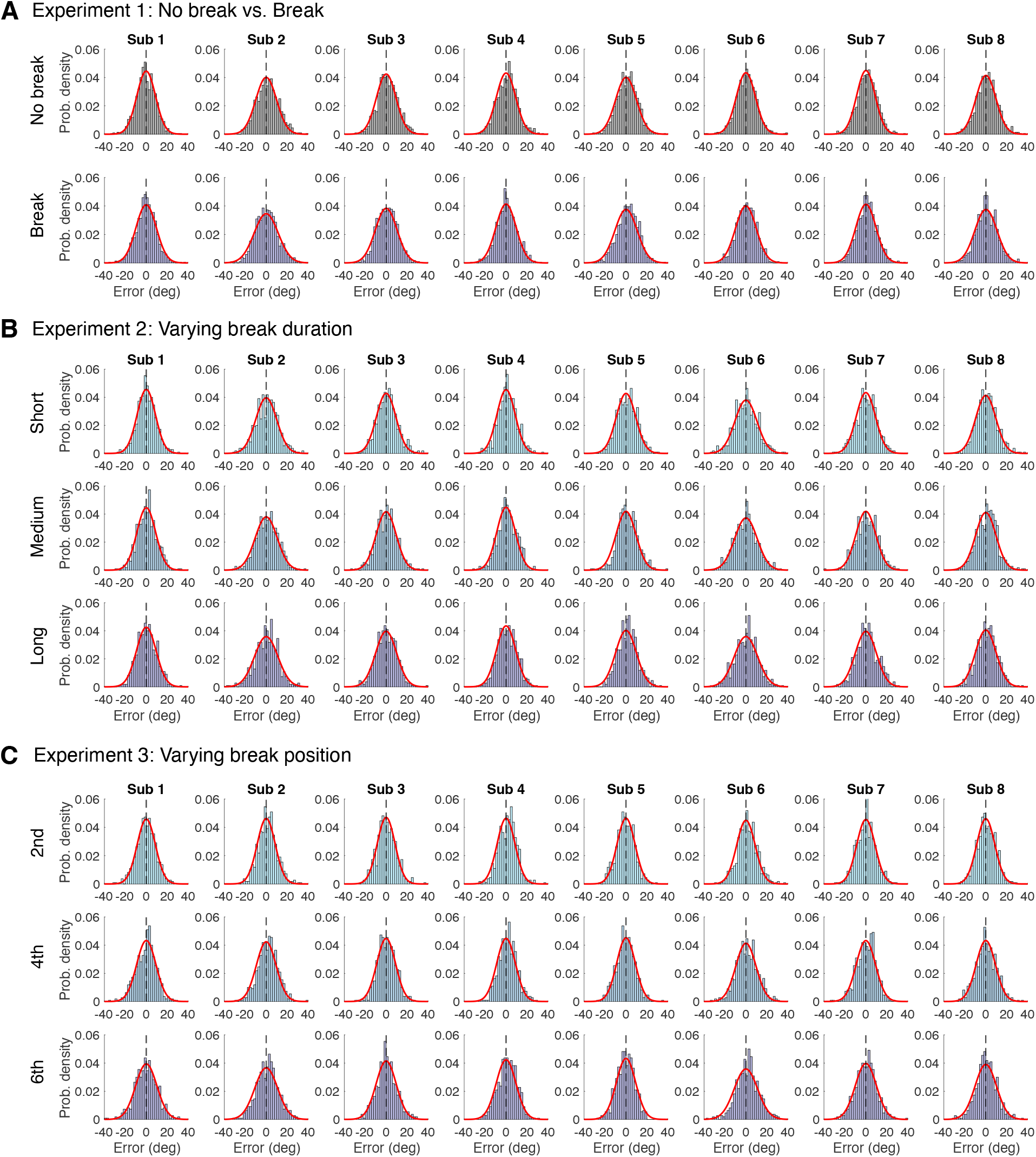
Error distributions of individual subjects’ responses relative to the true generative mean, together with the corresponding model fits (red line).

**Figure S3:**
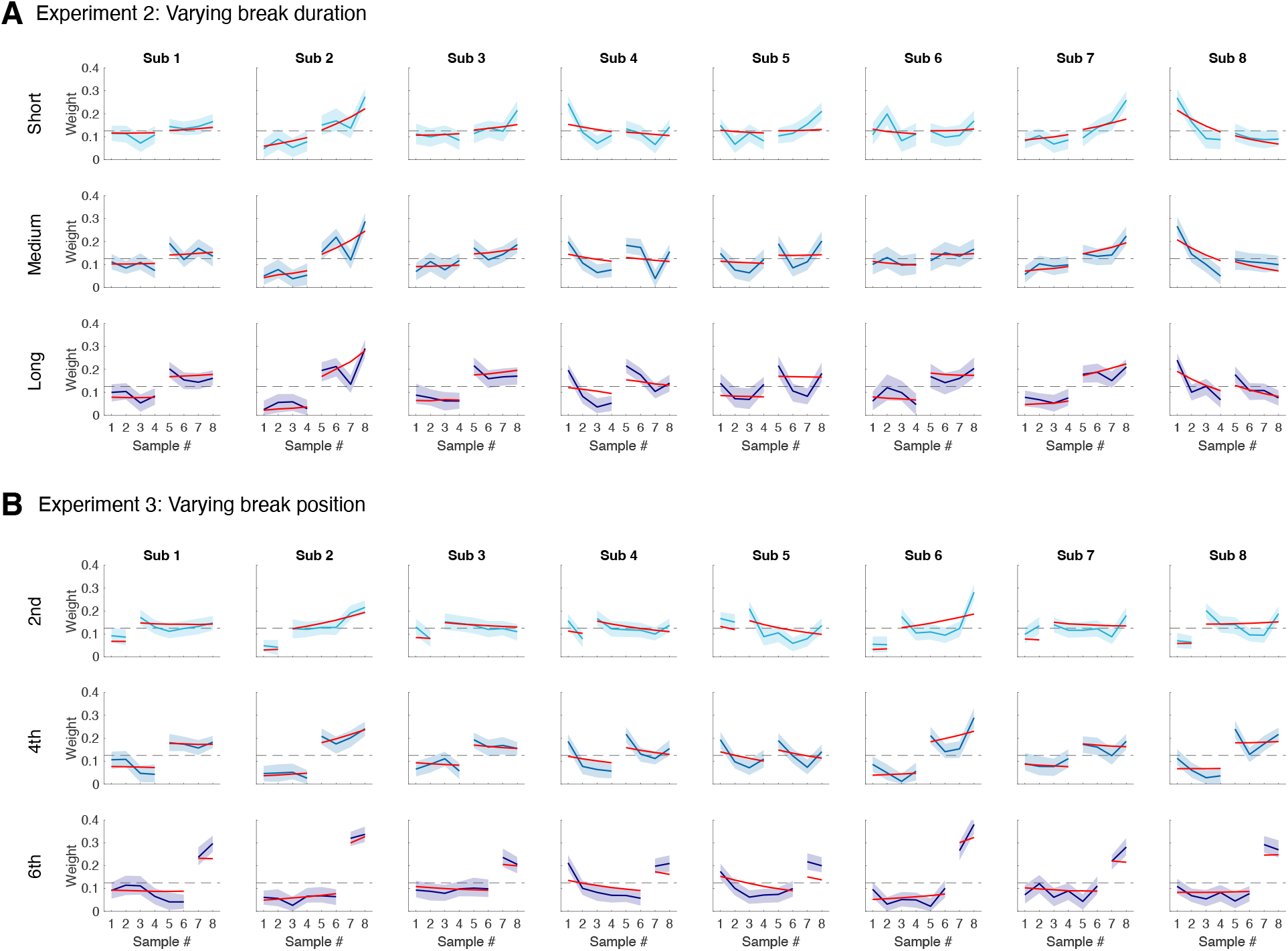
Circular regression weights of individual subjects in Experiment 2 (A) and Experiment 3 (B). Shaded areas indicate 95% confidence intervals based on 200 bootstrap samples of the data. Red curves indicate the model predicted weights based on joint model fits.

### Bump attractor network simulation of Experiment 1

**Figure S4:**
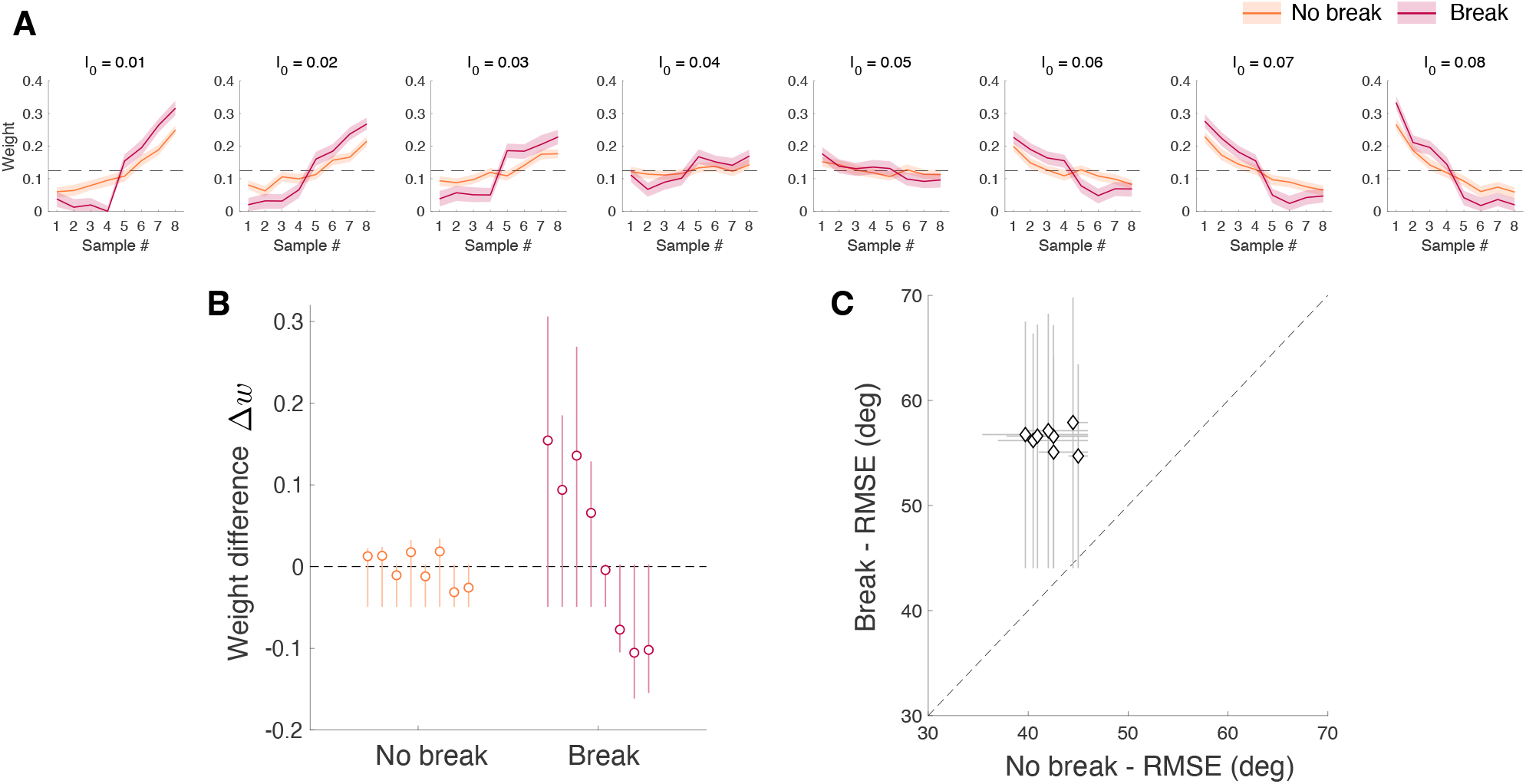
Bump attractor network simulations for “No break” and “Break” trials based on the model of Esnaola-Acebes, Roxin, and Wimmer (2022)^16^. (A) Circular regression weights computed from simulations for different values of the excitatory drive *I*_0_. (B) Weight difference Δ*w* between the 5th and 4th stimulus samples for “No break” and “Break” trials across different *I*_0_ values. (C) Estimation performance, measured as root mean squared error (RMSE), was significantly higher for “Break” trials across all tested values of *I*_0_.

